# Loss of Tip60-dependent H2A.Z acetylation reprograms cardiomyocyte identity toward a regenerative state

**DOI:** 10.1101/2024.01.11.575312

**Authors:** Xinrui Wang, Katherine R. Harty, Tina C. Wan, Zhuocheng Qu, John W. Lough, John A. Auchampach

## Abstract

Approaches to regenerate cardiomyocytes (CMs) after cardiac injury have been insufficient. Toward this end we are targeting the acetyltransferase Tip60, encoded by the *Kat5* gene, based on the rationale that its pleiotropic functions block CM proliferation at multiple checkpoints. We previously reported that genetic depletion or pharmacological inhibition of Tip60 in mice after myocardial infarction (post-MI) reduces scarring, restores function, and activates the cell-cycle in CMs, although it remains unresolved whether daughter CMs are generated. For pre-existing adult CMs in the infarcted adult heart to proliferate, they must first undergo dedifferentiation, a process characterized by loss of maturity, epithelial to mesenchymal transitioning (EMT), and a metabolic shift from fatty acid oxidation (FAO) to glycolysis. Recent studies indicate that Tip60 is required to maintain the differentiated state of hematopoietic stem cells and neurons via site-specific acetylation of the histone variant H2A.Z, specifically H2A.Zac^K^^4/K^^7^. Based on these findings we have examined H2A.Zac^K^^4/K7^ levels and the expression of dedifferentiation-associated markers in adult hearts following CM-specific knockout of Tip60. In infarcted/Tip60-depleted hearts, H2A.Zac^K4/K7^ was largely extinguished in CM nuclei, accompanied by the altered expression of genes consistent with EMT induction, extracellular matrix softening, and reduced FAO. Seahorse metabolic analyses of isolated CMs indicated that Tip60 depletion promotes an oxidative-to-glycolytic metabolic transition. In parallel, CUT&Tag analysis of nuclei isolated from heart tissue revealed that Tip60 depletion significantly reduced H2A.Zac^K4/K7^ occupancy within the promoter/transcription start sites of 47 genes that are highly CM-enriched, including cardiac maturity genes that are preceded in development by the expression of embryonic isoforms. RNAseq and RT-qPCR revealed that expression of these genes’ adult isoforms, as well as ∼50% of maturity genes that are not preceded by embryonic isoforms, were downregulated in Tip60-depleted hearts. These findings are consistent with the hypothesis that the Tip60 H2A.Zac^K4/K7^ axis maintains the differentiated state of CMs, constituting a major barrier to cardiac regeneration, justifying clinical targeting.

## INTRODUCTION

The burden of ischemic heart disease remains large, with myocardial infarction (MI) and heart failure accounting for ∼15% of deaths in the U. S.^1^ Because billions of cardiomyocytes (CMs) can be lost, the goal of protecting and re-muscularizing the myocardium constitutes a major clinical challenge. Mitigation of cardiac dysfunction caused by CM loss and replacement by scar tissue requires the regeneration of lost CMs, i.e. remuscularization. Unlike most organs, the adult heart cannot replace CMs due to their profound state of proliferative senescence that becomes established soon after birth.

With the goal of releasing surviving CMs from the non-proliferative state, recent research has targeted proteins that induce or inhibit CM proliferation via respective over-expression or knockout (KO) of genes in mouse models (reviewed in^2^). While these findings have shown that the CM cell-cycle can be re-activated, the extent of proliferation and functional improvement in most instances is modest, which is not surprising considering the array of inhibitors that prevent CMs from dividing. To disengage these inhibitors post-MI we are targeting the Tip60 acetyltransferase (*Kat5*), per a rationale based on findings in cultured cell-lines showing that Tip60 acetylates proteins that block the cell-cycle at three checkpoints: (i) G_1_/S, by maintaining p21 at inhibitory levels^3,4^, (ii) G_2_/M, by activating Atm and the DNA damage response (DDR)^5^, and (iii) cytokinesis, via acetylation of Aurkb^6^; also, because Tip60-mediated acetylation of p53 induces apoptosis^7–9^, its targeting may be cardioprotective. Consistent with these *in vitro* findings, our recent work using an *in vivo* Tip60 KO model has shown that genetic depletion of Tip60 from CMs post-MI restores cardiac function to near pre-MI baseline levels, with reduced scarring and cell-cycle activation^10,11^, and that these benefits are phenocopied by administering drugs that inhibit Tip60’s acetyltransferase activity^12,13^. However, it remains unclear whether activation of the CM cell-cycle results in bona fide proliferation, i.e. the generation of daughter CMs that are mononuclear and diploid.

It is now accepted that in order for mature CMs in the injured adult heart to resume proliferation, they must undergo dedifferentiation^14–20^, a process wherein the expression of mature adult cardiac genes becomes diminished as embryonic genes are re-expressed (reviews^21–24^). CM dedifferentiation is highlighted by cell-cycle activation^14–16^, an epithelial-to-mesenchymal-like transition (EMT)^17^, extracellular matrix (ECM) softening^16,18,25–27^, and as recently reported, inhibition of fatty acid oxidation (FAO)^19^. Here, we report that post-MI depletion of Tip60 in CMs is accompanied by the near-total loss of lysine K4 and K7 acetylation in the histone variant H2A.Z (H2A.Zac^K^^4/K7^), a Tip60-mediated epigenetic modification recently shown to be required to maintain the differentiated state of hematopoietic stem cells (HSCs)^28^ and neurons^29^. This was accompanied by CM dedifferentiation as evidenced by differential gene expression indicating cell-cycle activation, EMT, ECM softening, and reversion from adult to embryonic metabolism as verified by Seahorse assay. Using CUT&Tag, we also show that H2A.Zac^K4/K7^ is associated with genetic motifs and GO terms respectively associated with CM transcription factor binding and muscle development/differentiation, and that Tip60 KO reduces H2A.Zac^K4/K7^ levels in promoter/TSS loci of genes considered to be markers of cardiac maturity. Diminished H2A.Zac^K4/K7^ levels correlated with reduced transcription in approximately 50% of genes associated with CM maturation, most remarkably those preceded in development by the expression of embryonic isoforms. These results, in combination with our previous findings, support the theory that Tip60’s multiple targets collaborate to block CM regeneration, justifying its pharmaceutical targeting after myocardial infarction.

## MATERIALS & METHODS

### Heart Tissue Processing

These experiments were performed on samples of left ventricular heart tissue obtained from ∼12-week-old naïve/Tip60-depleted mice, and from the remote zone of infarcted/Tip60-depleted mice, as previously described^11–13^. All protocols adhered to the National Institutes of Health (NIH) Guide for the Care and Use of Laboratory Animals (NIH Pub. Nos. 85-23, Revised 1996) as described in Animal Use Application #000225 and approved by the Medical College of Wisconsin (MCW) Institutional Animal Care and Use Committee (IACUC). MCW has welfare assurance status from the Office of Laboratory Animal Welfare (A3102-01). Mice were on a mixed B6/sv129 genetic background. Experimental groups contained similar numbers of males and females. Genotyping was performed by PCR as previously described^11^ using primer pairs listed in Supplemental Table 1. Tissue samples for RNA and protein isolation were respectively placed in TRIzol (Thermo-Fisher #15596026), and Histone Extraction Buffer (Active Motif #37513) containing Halt’s protease/phosphatase inhibitor cocktail (Thermo-Fisher #78440) with deacetylase inhibitors (MedChemExpress #HY-K0030), minced, homogenized with a teflon pestle, and stored at -80° C until further processing.

### RT-qPCR

Heart tissue was homogenized using a Kimble #749540 Teflon pestle and stored in TRIzol at -80° C. Upon thawing, RNA was purified via PureLink RNA Mini-Kit (ThermoFisher #12183018A) including the DNA removal step (Thermo-Fisher #12185-010) per manufacturer’s instructions. RNA yield & quality were determined via 260/280 using an Eppendorf Biophotometer Plus instrument. After diluting RNA to 1.0 µg/14 µl NFDW and adding 4.0 µl 5x VILO reaction mixture (ThermoFisher #11754050), cDNA synthesis was initiated with 2.0 µl 10x SuperScript Enzyme Mix (ThermoFisher #11754050) and samples were incubated in an Applied Biosystems Veriti 96-well Thermocycler programmed as follows: 10 minutes/25°C, 60 minutes/42°C, 5 minutes/85°C. Synthesized cDNA was diluted in NFDW to ∼3 ng/µl and stored at -20° C. For qPCR amplification, each biological replicate (i.e., each heart) was subjected to three technical triplicates carried out in a total volume of 10 µl using 384-well arrays, with each well containing 1x Taqman Fast-Advanced Master Mix (ThermoFisher #4444557), 1x Taqman Probe Kit (Supp. Table 2), and 12.5 ng cDNA as template. cDNA was amplified as previously described^11^ with results processed using Bio-Rad CFX Manager 3.1 software.

### Western Blotting

Histones were purified using Active Motif Histone Purification Mini Kit #40026 per the manufacturer’s recommendations. All procedures were performed at 0-4 °C using a minimal volume of low pH Extraction Buffer containing protease and deacetylase inhibitors. Tissue (∼50 mg) dispersed by Dounce homogenization and microtip sonication was centrifuged (5 min/21,000g), after which supernatants were adjusted to pH 8.0 with 5X Neutralization Buffer. Crude histones were de-salted in a Zeba Spin Desalting Column (Thermo-Fisher #89883) followed by addition of sample buffer (Cell Signaling Technology #12957). For electrophoresis, samples in pre-cast Bio-Rad 4-20% acrylamide gels were separated at 100 V, followed by transfer (60 min/100 V) to 0.2 μm nitrocellulose (Thermo-Fisher #88013). Blots were blocked using 5% non-fat dry milk/20 mM Tris (pH 7.6)/150 mM NaCl/0.1%Tween-20 (5% NFDM/TBST), or 5% BSA in TBST. Primary antibodies were anti-acetylated histone H2A.Z (Lys4/Lys7, 1:1000, Cell Signaling Technology #75336) and anti-bulk histone H2A.Z (1:1000, Active Motif #39013), followed by secondary antibody application (HRP-linked goat antirabbit (1:2000, Cell Signaling Technology #7074). Blots were reacted with primary antibody in 5% NFDM/TBST or 5% BSA/TBST overnight at 4°C. The next day, secondary antibodies were diluted in 5% BSA/TBST and applied for 60 minutes at RT. Reacted blots were covered with Pierce™ ECL horseradish peroxidase substrate (ThermoFisher #32209; 1 min/RT) followed by chemiluminescent imaging and densitometry, respectively using Bio-Rad ChemiDoc and ImageJ software.

### Immunostaining

Mice were injected with 1 mg 5’-bromo-2’-deoxyuridine (BrdU; Sigma #B9285) ∼24 hours before harvest. Following removal, hearts were perfused with ice-cold cardioplegic solution, atria were removed, and ventricles were fixed overnight in fresh ice-cold 4% paraformaldehyde/PBS followed by processing through ethanol series and embedment in paraffin. Sections (4 µm) were mounted on microscope slides, dewaxed, and subjected to antigen retrieval (20 min at 100°C in 10 mM trisodium citrate pH6.0/0.05% Tween-20) followed by 30 min cooling at RT, and blocked with 2% goat or 5% donkey serum/0.1% Triton-X-100 in PBS. Primary antibodies against acetyl-histone H2A.Z (Lys4/Lys7, 1:100, Cell Signaling Technology #75336), histone H2A.Z (1:200, Active Motif #39013), Nkx2.5 (1:250, Abcam #ab106923), cardiac Troponin-T (cTnT, 1:2000, Abcam #ab8295), α-smooth muscle actin (1:100, Dako #M0851), and H3K4^me3^ (1:600, Active Motif #39160) were diluted in blocking buffer and applied overnight at 4°C. Secondary antibodies were goat antirabbit 568 (1:500, Invitrogen #A-11036), goat anti-mouse 488 (1:500, Invitrogen #A-11029), donkey anti-rabbit 555 (1:500, Invitrogen #A-31572), and donkey anti-goat 488 (1:500, Invitrogen #A-32814), which were applied for one hour in the dark.

Microscopy utilized a Nikon Eclipse 50i microscope equipped with a Nikon DSU3 digital camera. For counting, at least 300 CMs were evaluated in six random 400x fields. CM identity was verified by immunostaining cTnT and the nuclear marker Nkx2.5. Percentages of CMs exhibiting acetylated and bulk histone H2A.Z were assessed by monitoring signal in the Texas Red channel after confirming CM identity in the FITC channel. To identify a nucleus as positive for acetylated or bulk histone H2A.Z, only nuclei that were at least half-filled with fluorescence were counted. In infarcted hearts, acetylated and bulk histone H2A.Z immunoreactivity was assessed in both the remote zone, i.e., the area of myocardium ∼2 mm distal to the infarct boundary, and in the border zone, the area adjacent to the infarct zone. WGA staining was performed using Thermo-Fisher #W11263 Alexa Fluor 350 conjugate.

### RNAseq

RNA quality was determined using an Agilent BioAnalyzer 2100. RNA libraries were prepared by BGI Americas. Samples were tested for strand-specific mRNA sequencing on a DNBSEQ platform. SOAPnuke v1.5.2 was used to remove reads possessing N content >5%, reads containing adaptor, and reads with low base quality (<15). Filtered clean reads (>45 M per sample) were stored in FASTQ format and aligned to the *Mus musculus* genome (GCF 000001635.26 GRCm38.p6) using HISAT2 v 2 0.4. Bowtie2 was used to align clean reads to the reference genes. A total of 17,714 genes were detected. The average alignment ratio of the sample comparison genome was 96.63%. Gene expression levels were calculated by RSEM (v1.2.28) and quantified by fragments per kilobase of transcript per million reads mapped (FPKM). DEGs for each experimental group, as compared to control, were detected using DESeq2. The phyper function in R software was used for enrichment analysis and calculation of P values; Q values were obtained by correction of the P value.

### Seahorse Metabolic Assay

Mitochondrial oxidation and glycolysis were respectively evaluated by analyzing oxygen consumption rate (OCR, pmol/min) and extracellular acidification rate (ECAR, mpH/min) using a XF96 Extracellular Flux Analyzer (Agilent). On the day before assay, 200 μL of XF Calibrant were loaded into each well of a 96-well utility plate included with the sensor cartridge, and the sensors were submerged in a 37 °C non-CO_2_ incubator overnight. On the day of assay, primary adult CMs were isolated and plated in wells coated with laminin (20 μg/ml) at 2.5 x 10^3^ cells/well for mitochondrial oxidation assays, or at 5 x 10^3^ cells/well for glycolysis assays; medium consisted of MEM supplemented with 10% FBS, 2 mM glutamine, 100 U/ml penicillin, 100 μg/ml streptomycin, 2 mM ATP, and 15 μM blebbistatin. One hour prior to assay, the cell culture medium was changed to assay medium containing non-buffered XF RPMI (phenol red-free; Agilent) supplemented with 1mM pyruvate and 2mM L-glutamine and 10mM glucose for the mitochondrial oxidation assay or 2mM Lglutamine for glycolysis assay.

For the mitochondrial oxidation assay, OCRs were obtained from the slope of change in oxygen over time. Measurement of baseline OCR was followed by sequential injection of substrates/inhibitors, to final concentrations of 2 μM oligomycin (ATP synthase inhibitor), 1 μM carbonyl cyanide p (trifluoromethoxy) phenylhydrazone (FCCP, uncoupler of oxidative phosphorylation in mitochondria), and 1 μM rotenone and antimycin A (electron transport chain complex I and III blocker). OCR parameters assessed were (1) Basal Respiration, representing the last rate measurement before oligomycin injection minus the non-mitochondrial respiration rate (the minimum rate measurement after antimycin A injection), (2) Maximal Respiration, the maximum rate measurement after FCCP injection (minus non-mitochondrial respiration rate), (3) Spare Respiratory Capacity: the maximum respiration minus Basal Respiration, and (4) proton leak: minimum rate measurement after Oligomycin injection.

For the glycolysis assay, ECARs were obtained from the slope of change in H^+^ concentration over time. After assessing baseline ECAR, ECARs were analyzed after sequential injection of the following substrates/inhibitors to final concentrations of 20 mM glucose, 2 µM oligomycin, and 50 mM 2-DG. Each ECAR parameter was calculated as: (1) Glycolysis, the maximum rate measurement before oligomycin injection minus the last rate measurement before glucose injection; (2) Glycolytic Capacity, the maximum rate measurement after oligomycin injection minus the last rate measurement before glucose injection, and (3) NonGlycolytic Acidification, the last rate measurement prior to glucose injection. Each cycle consisted of a 1.5-minute mixing, a 2-minute waiting, and a 1.5-minute measuring phase. All calculated OCR and ECAR parameters were normalized to DNA content (DAPI intensity).

### Cleavage Under Targets & Tagmentation (CUT&Tag) Assay

H2A.Zac^K4/K7^ peaks were assessed using H2A.Zac^K4/K7^ antibody (Cell Signaling Technology #75336), which was qualified for CUT&Tag by comparing genomic targets with those recognized by a pre-qualified antibody that specifies bulk H2A.Z (Active Motif #39113). As shown in Supp. Fig. 1, (i) anti-H2A.Zac^K4/K7^ specifically recognized histone H2A.Z (Supp. Fig. 1a), (ii) 15-23% of peaks were in promoter/TSS loci (Supp. Fig. 1b), (iii) targets were enriched in cardiac transcription factor binding motifs (Supp. Fig. 1c), and (iv) were associated with Gene Ontology (GO) terms denoting cardiac development and differentiation (Supp. Fig. 1d). Procedurally, ∼10 mg left ventricular tissue from hearts of 14-week-old naïve adult mice were processed for CUT&Tag by Active Motif (Carlsbad, CAusing tissue kit #53170. One million nuclei from each heart were attached to Concanavalin A beads, followed by overnight incubation with anti-H2A.Zac^K4/K7^. After attachment of guinea pig secondary antibody, bound targets were subjected to cleavage and tagmentation via incubation with protein-A-Tn5 transposase/sequencing adapters, followed by DNA purification, PCR amplification using i7/i5 primer pairs selective for individual samples, and isolation on SPRI beads. DNA libraries were sequenced on Illumina NextSeq 550 (8 million reads, 38 pairedend). For details see instructions in Active Motif CUT&Tag-IT kit #53170. Fraction of Reads In Peaks (FRIP) percentages ranged from 36.8 to 43.9, with total peak numbers ranging from 34,810 to 53,327. Peak annotation and motif analysis were assessed using HOMER^30^. Across all samples, upset plotting revealed a consensus set of ∼30,000 peaks. H2A.Zac^K4/K7^ peaks were imaged using the WashU Epigenome Browser^31^ (https://epigenomegateway.wustl.edu/browser).

### Statistics

DEG heatmaps were generated by the Dr.Tom platform (BGI Americas) according to DESeq2 analysis results. GO enrichment analysis of annotated DEGs was performed by Phyper based on Hypergeometric test. To minimize false-positives, the identification of statistically significant differences between H2A.Zac^K4/K7^ peaks by CUT&Tag was assessed using adjusted p-value (i.e., p-adjust) except as otherwise noted. For RNAseq, significant terms and pathways were corrected by Q ≤0.05. All determinations were performed in blinded fashion and reported as means (±SEM), compared using unpaired two-tailed Student t testing; p<0.05 was considered statistically significant.

## RESULTS

### Genetic Depletion of Tip60 in CMs Abolishes Acetylation of Histone H2A.Z

These experiments compare 12-week-old adult control (denoted *Kat5^f/f^*, i.e., *Kat5^flox/flox^*) with experimental (*Kat5*^Δ^*^/^*^Δ^, i.e., *Kat5^flox/flox;Myh6-merCremer^*) mice in which the *Kat5* gene encoding Tip60 was disrupted by injecting tamoxifen for three consecutive days to activate *Myh6*-driven Cre-recombinase (hereafter, ‘Cre’). In infarcted hearts treated with tamoxifen beginning on the third day post-MI (hereafter ‘infarcted/Tip60-depleted’ hearts), expression of *Kat5* in samples from whole heart tissue was reduced ∼25% (Supp. Fig. 2d); although depletion was presumably greater in CMs *per se*, this extent of reduction was considered modest. By contrast, in control (*Kat5^f/f^*) hearts employed to assess potential off-target effects of Cre, expression of *Kat5* was unaffected (Supp. Fig. 2d), and effects on other genes were minimal (88 DEGs; Supp. Fig. 2b,c).

Using this model, we recently reported that depletion of Tip60 post-MI preserves cardiac function, reduces scarring, and activates the cell-cycle in CMs of neonatal^10^ as well as adult^11^ hearts. Here, as shown by immunostaining (Fig. 1a), depletion of Tip60 resulted in the absence of H2A.Zac^K4/K7^ signal in nearly all CM nuclei. This is the most pronounced effect of Tip60 depletion we have observed in this as well as previous studies^10,11,13^. To verify that H2A.Zac^K4/K7^ depletion was confined to nuclei in CMs, cellular identity was confirmed using the nuclear marker Nkx2-5 (Supp. Fig. 3). By contrast, using an antibody that detects bulk (i.e., both acetylated and non-acetylated) H2A.Z, no effects of Tip60 depletion were seen (Supp. Fig. 4). Controls indicated that CM-specific depletion of H2A.Zac^K4/K7^ did not represent an off-target effect of Cre (Supp. Fig. 5). Depletion of H2A.Zac^K4/K7^ was verified by western blotting (Fig. 1b); despite blotting protein from whole heart tissue in which most cells are non-CMs, levels of H2A.Zac^K4/K7^ were reduced by ∼two-thirds, while levels of bulk H2A.Z were modestly increased (Supp. Fig. 6).

**Figure 1.**
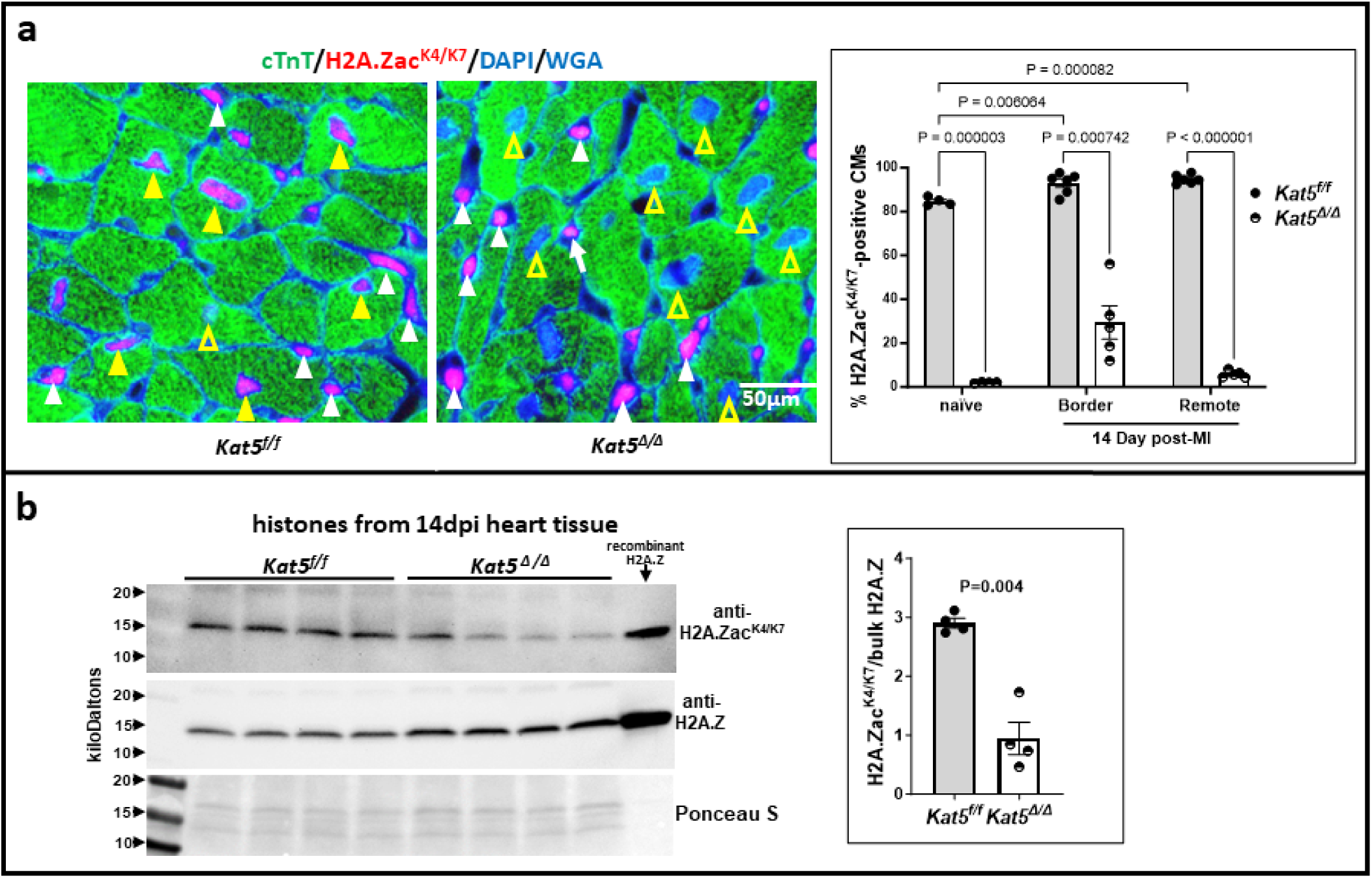
H2A.Zac^K4/K7^ depletion in Tip60-depleted CMs. **Panels a-b** respectively show representative images of infarcted adult control (*Kat5^f/f^*) and infarcted/Tip60-depleted (*Kat5*^Δ^*^/^*^Δ^) hearts. In the latter, Tip60 was depleted from CMs by injecting tamoxifen for three days beginning on day 3 post-MI. Cardiac tissue was processed for immunostaining (**a**) and western blotting (**b**) on day 14 post-MI. **a, Immunostains:** In *Kat5^f/f^* control hearts, acetylated H2A.Z (i.e., H2A.Zac^K4/K7^; red fluorescence) is present in nuclei of most non-CMs (white arrowheads) and CMs (filled yellow arrowheads). In Tip60-depleted hearts (*Kat5*^Δ^*^/^*^Δ^), H2A.Zac^K4/K7^ is present in non-CM nuclei (white arrowheads), but is conspicuously absent in most CM nuclei (open yellow arrowheads). At right, percentages of H2A.Zac^K4/K7^-positive CMs were enumerated by blinded observers. **b, Western blots** showing acetylated (upper) vs. non-acetylated (i.e., bulk, middle) H2A.Z protein in heart tissue. Right, densitometry showing altered levels of H2A.Zac^K4/K7^ normalized to bulk H2A.Z. In **panels a & b**, *P<0.05 vs. *Kat5^f/f^*per unpaired Student’s t-test; means ±SEM. Each point represents a biological replicate (i.e., an individual mouse heart).

**Figure 2.**
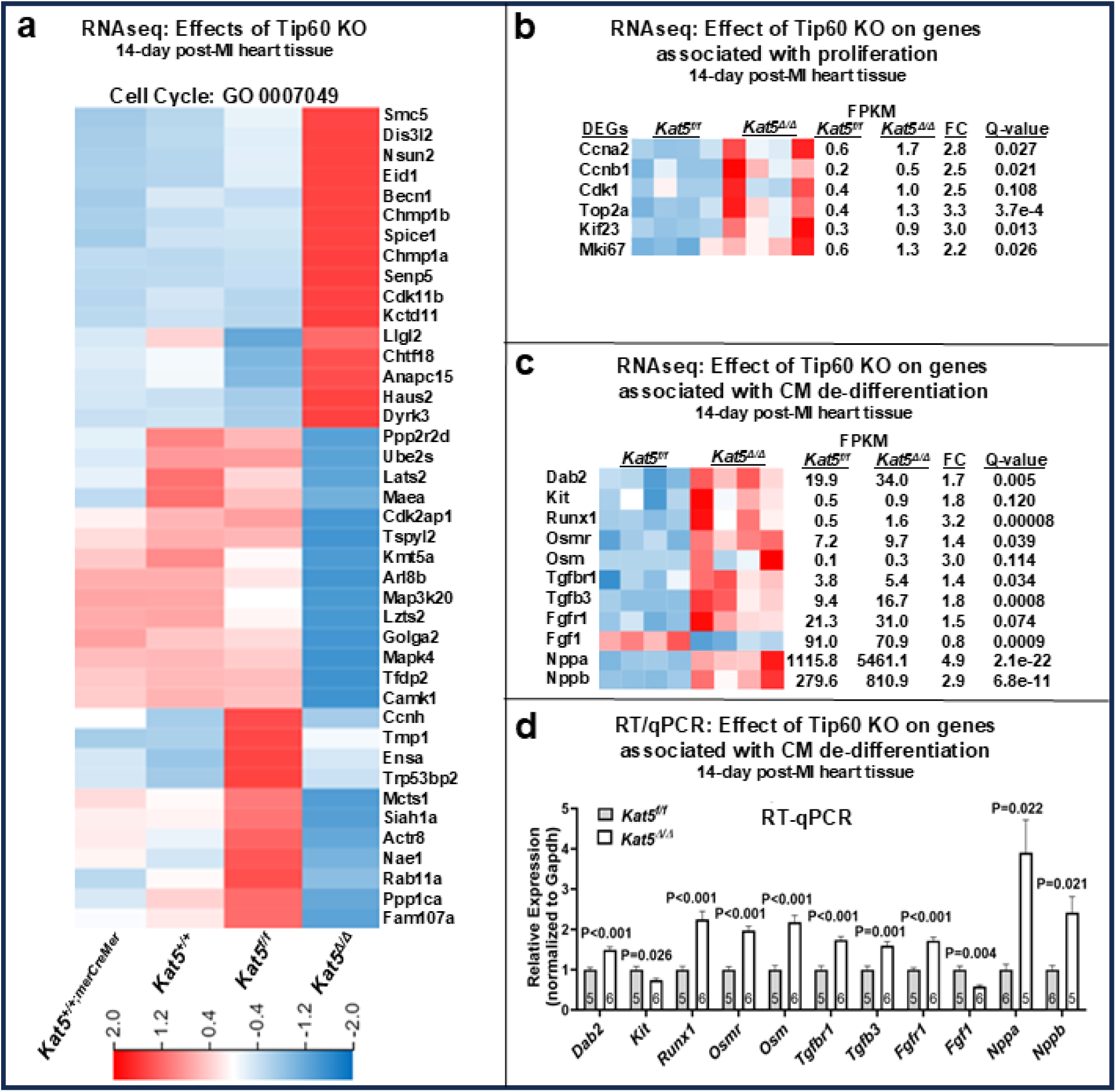
Expression of genes associated with dedifferentiation and proliferation signaling in infarcted/Tip60-depleted hearts at 14 days post-MI. **Panel a**: RNAseq results of Gene Ontology term ‘Cell Cycle’. **Panel b**: RNAseq data showing increased expression of pro-proliferative genes. **Panels c-d**: RNAseq (**c**) and verifying RT-qPCR (**d**) data showing increased expression of genes previously associated with CM dedifferentiation. *Kat5^f/f^* = control, *Kat5*^Δ*/*Δ^ = Tip60-depleted. In **panel a**, lanes labeled *Kat5^+/+^* and *Kat5^+/+;merCremer^* are controls to monitor off-target effects of Cre. In **b-c** each column represents a biological replicate. In **d**, each biological replicate (N, inside bar) is a single heart subjected to three technical replicates. Data are Mean ±SEM compared by unpaired, two-tailed Student’s t test with Welch’s correction.

**Figure 3.**
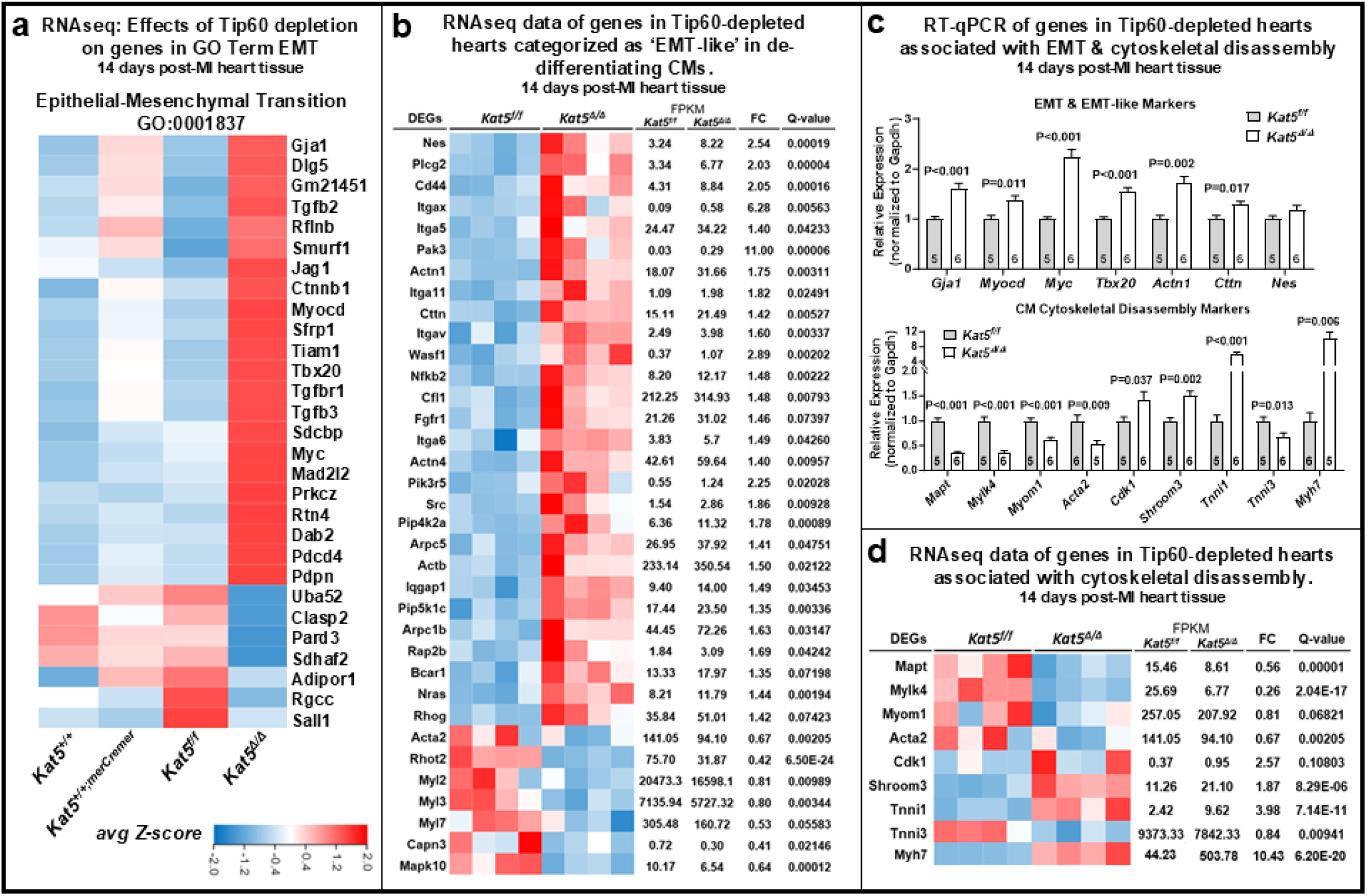
Expression of genes associated with epithelial-mesenchymal transition (EMT) & cytoskeleton stability in infarcted/Tip60-depleted hearts. **Panel a,** RNAseq data showing effect of Tip60 on genes in the Gene Ontology term ‘Epithelial-Mesenchymal Transition’ (EMT). **Panel b**, RNAseq data showing misexpressed EMT-like genes. **Panel c**, RT-qPCR verification of mis-expressed EMT and EMT-like (upper) and cytoskeletal (lower) genes. **Panel d**, RNAseq data showing mis-expressed cytoskeletal genes. *Kat5^f/f^* = control, *Kat5*^Δ*/*Δ^ = Tip60-depleted. In panel **a**, *Kat5^+/+^*vs. *Kat5^+/+;merCremer^* assess off-target effects of Cre. In **b and d**, each column represents a biological replicate. In **c**, each biological replicate (N) is a single heart subjected to three technical replicates. RT-qPCR data are Mean ±SEM compared by unpaired two-tailed Student’s t tests with Welch’s correction.

**Figure 4.**
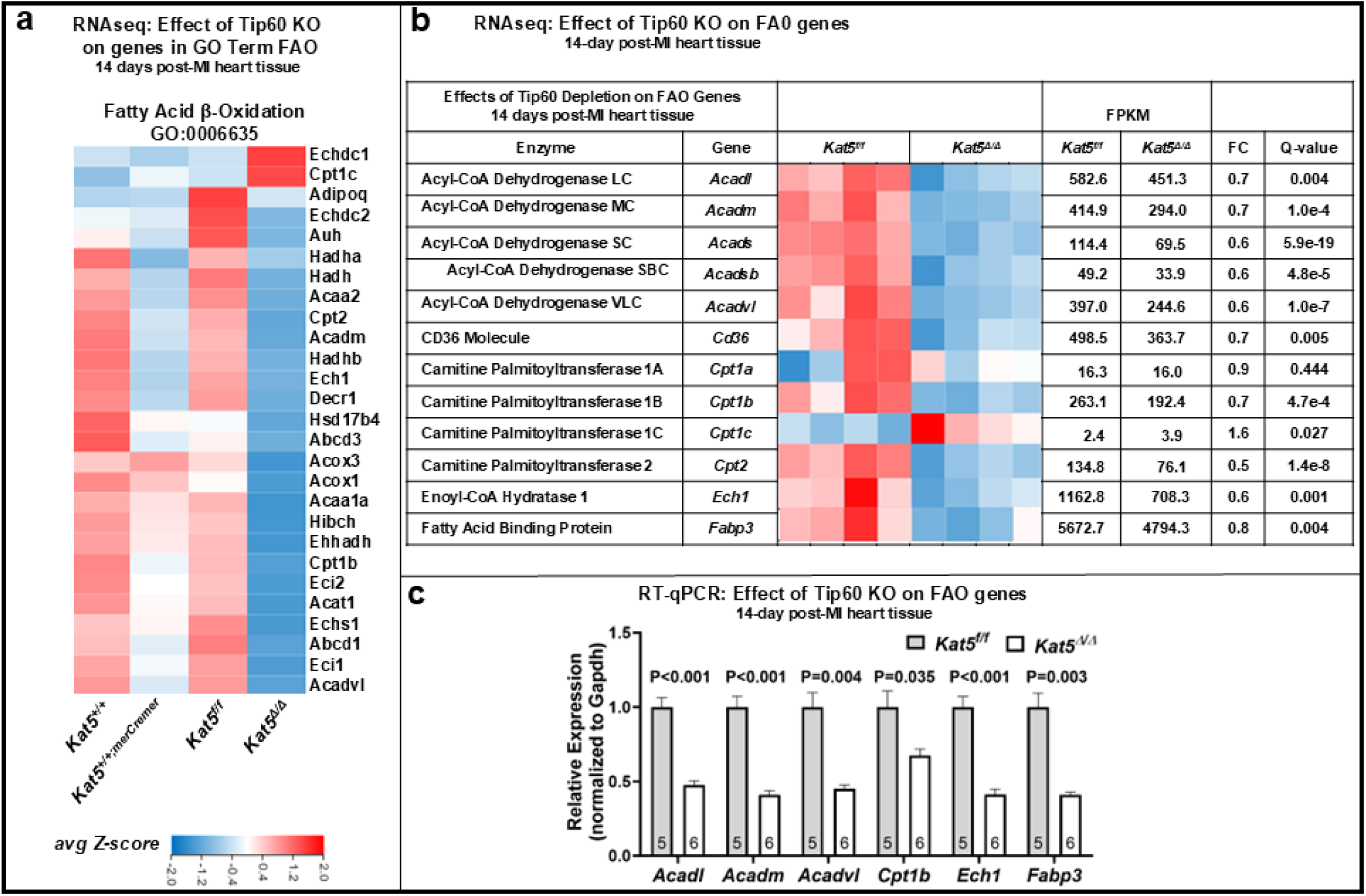
Expression of genes associated with fatty acid oxidation in infarcted/Tip60-depleted hearts. **Panel a,** RNAseq data showing mis-expressed genes in the GO term ‘Fatty Acid β-Oxidation’ **Panel b**, RNAseq data showing mis-expression of selected FAO genes in infarcted/Tip60-depleted hearts. Each column in the heatmap represents a biological replicate. **Panel c**, RT-qPCR verification of mis-expressed FAO genes. Each biological replicate (N) represents a single heart subjected to three technical replicates. Data are Mean ± SEM compared by unpaired, two-tailed Student’s t tests with Welch’s correction. *Kat5^f/f^* = control, *Kat5*^Δ^*^/^*^Δ^ = Tip60-depleted. *Kat5^+/+^* and *Kat5^+/+;merCremer^* assess off-target effects of Cre.

**Figure 5.**
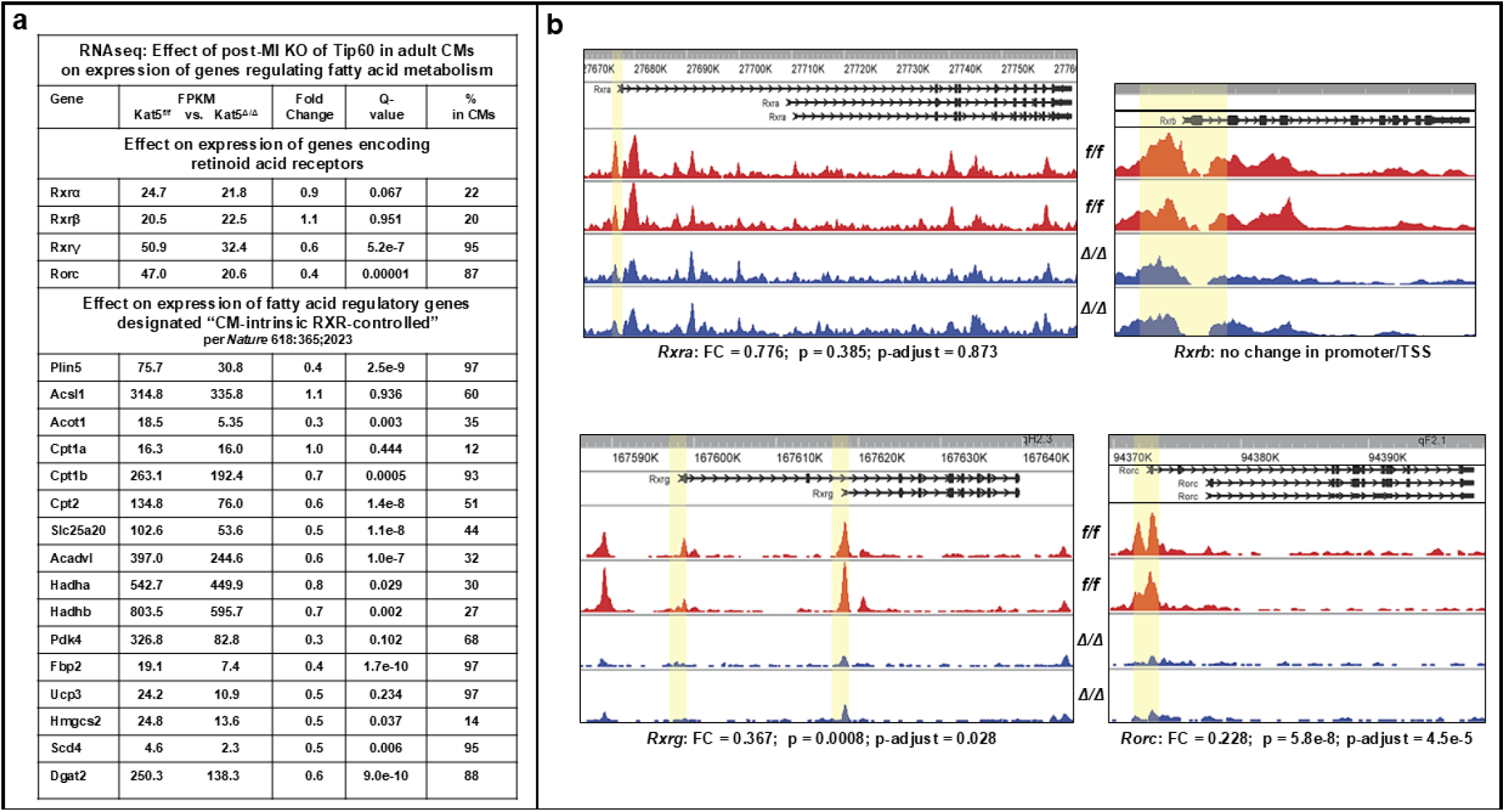
H2A.Zac^K4/K7^ depletion and diminished expression of genes encoding retinoic acid receptors *Rxrg* and *Rorc* and their targets in infarcted/Tip60-depleted hearts. **Panel a:** RNAseq data showing reduced expression of *Rxrg* and *Rorc* transcription factor genes (above) and of CM-intrinsic Rxr-controlled targets (below). The column labeled ‘% in CMs’ denotes enrichment of each RNA in CMs, relative to all cell types in heart tissue (per Tabula Muris). FPKM avgs based on Ns=5. **Panel b**, CUT&Tag genome browser images of Tip60-depleted adult hearts showing reduced levels of H2A.Zac^K4/K7^ in the promoter/TSS loci of genes encoding Rxrg and Rorc. f/f = control; Δ/Δ = Tip60 knockout.

**Figure 6.**
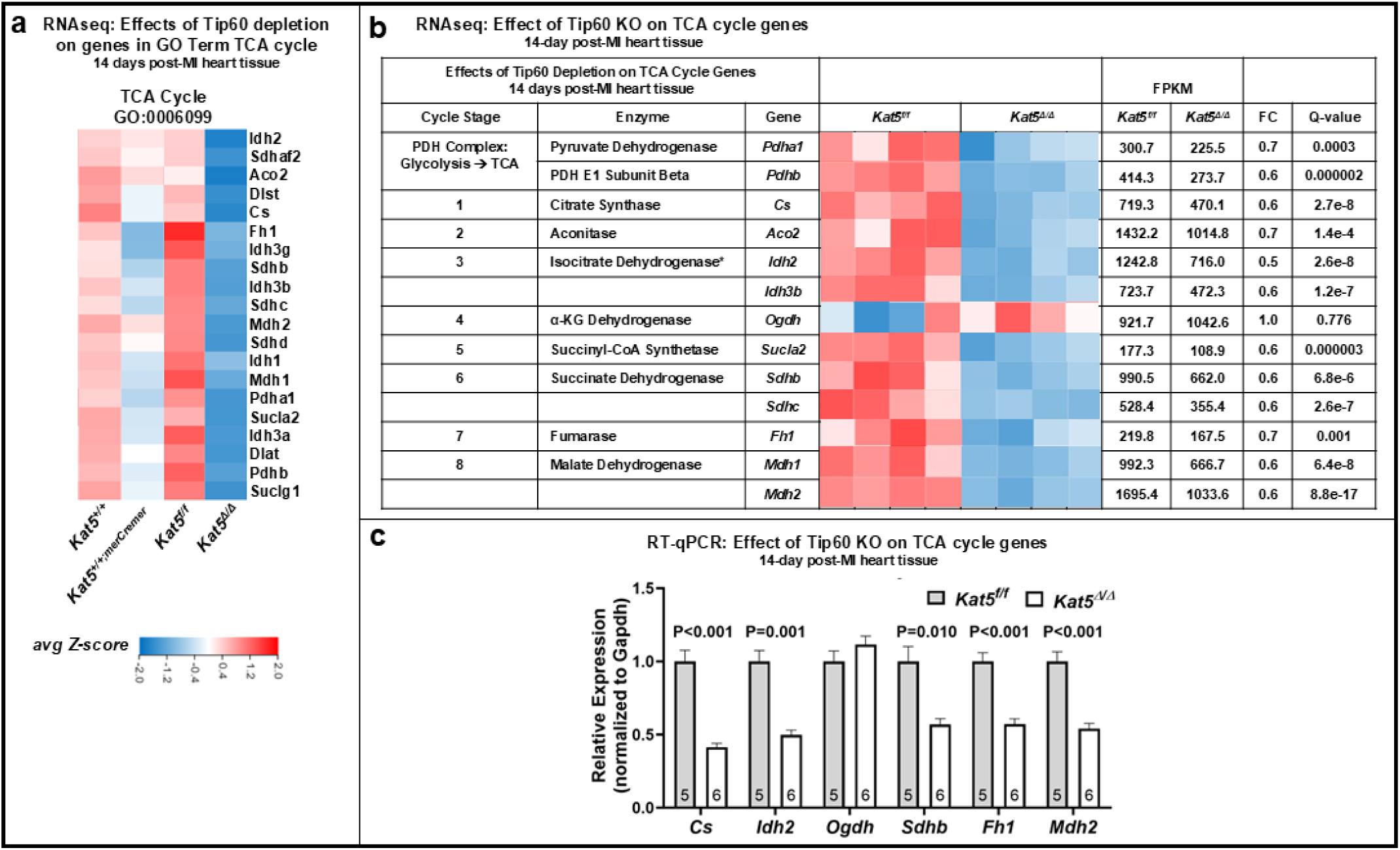
Expression of genes associated with GO term TCA cycle in infarcted/Tip60-depleted hearts. **Panel a**, RNAseq data showing mis-expressions of genes in the GO term ‘TCA cycle’. *Kat5^f/f^* = control, *Kat5*^Δ*/*Δ^ = Tip60-depleted. *Kat5^+/+^* and *Kat5^+/+;merCremer^* assess off-target effects of Cre. **Panel b**, RNAseq data showing mis-expressed genes that regulate TCA cycle stages in infarcted/Tip60-depleted hearts. Each column in the heatmap represents a biological replicate. **Panel c**, RT-qPCR verification of mis-expressed TCA cycle genes in infarcted/Tip60-depleted hearts. Each biological replicate (N; shown inside bar) is from a single heart, subjected to three technical replicates. Data are Mean ± SEM compared by unpaired, two-tailed Student’s t tests with Welch’s correction.

### Genome-Wide Impact of Depleting Tip60 in CMs on Gene Expression

Effects of depleting Tip60 in CMs on gene expression in adult hearts were assessed by RNAseq. As shown in Supplemental Figure 2, tamoxifen-mediated ablation of the *Kat5* gene in infarcted hearts resulted in the differential expression of 4,049 genes (DEGs, Supp. Fig. 2c), among which the top 10 up-regulated GO terms included cell adhesion, migration, cycling, and division, and the top 10 down-regulated GO terms included fatty acid metabolic, β-oxidation, and TCA cycle processes (Supp. Fig. 2a). These data indicate that depletion of Tip60 in CMs markedly affects gene expression, in particular genes that regulate processes associated with differentiation and proliferation.

### Infarcted/Tip60-depleted Hearts Exhibit Increased Expression of Genes Associated with CM Dedifferentiation & Cell-Cycle Activation

It is generally accepted that adult CMs must dedifferentiate in order to permit the resumption of cell-cycling and regeneration after cardiac injury^15–17,20,32^. Based on findings that site-specific acetylation of H2A.Z by Tip60 is required to differentiate HSCs^28^ and neurons^29^, we focused on assessing how Tip60 depletion affects the expression of specific genes that have been associated with CM dedifferentiation and proliferation. RNAseq (Fig. 2c) and corroborating RT-qPCR (Fig. 2d) data showed that Tip60 depletion was associated with up-regulation of the genes encoding Dab2, and Runx1, previously shown to be induced by oncostatin-M (Osm) at the onset of CM dedifferentiation^14,15^; accordingly, increased expression of genes encoding Osm as well as its receptor (Osmr) was also noted. *Fgfr1* and *Tgfbr1*, which have also been associated with CM dedifferentiation^16^, as well as *Tgfb3*, were also increased in Tip60-depleted hearts (Fig. 2c-d). However *Fgf1*, which among members of the large Fgf family is selectively expressed in CMs^33^, was decreased (Figs. 2c-d; Supp. Fig. 8a). Although controls assessing off-target effects of Cre on wild-type hearts (Fig. 2a) indicated a trend toward increased expression of genes that were reduced by Tip60 KO, none attained statistical significance; also, Cre had no effect on any of the genes evaluated in Figure 2b-d.

**Figure 7.**
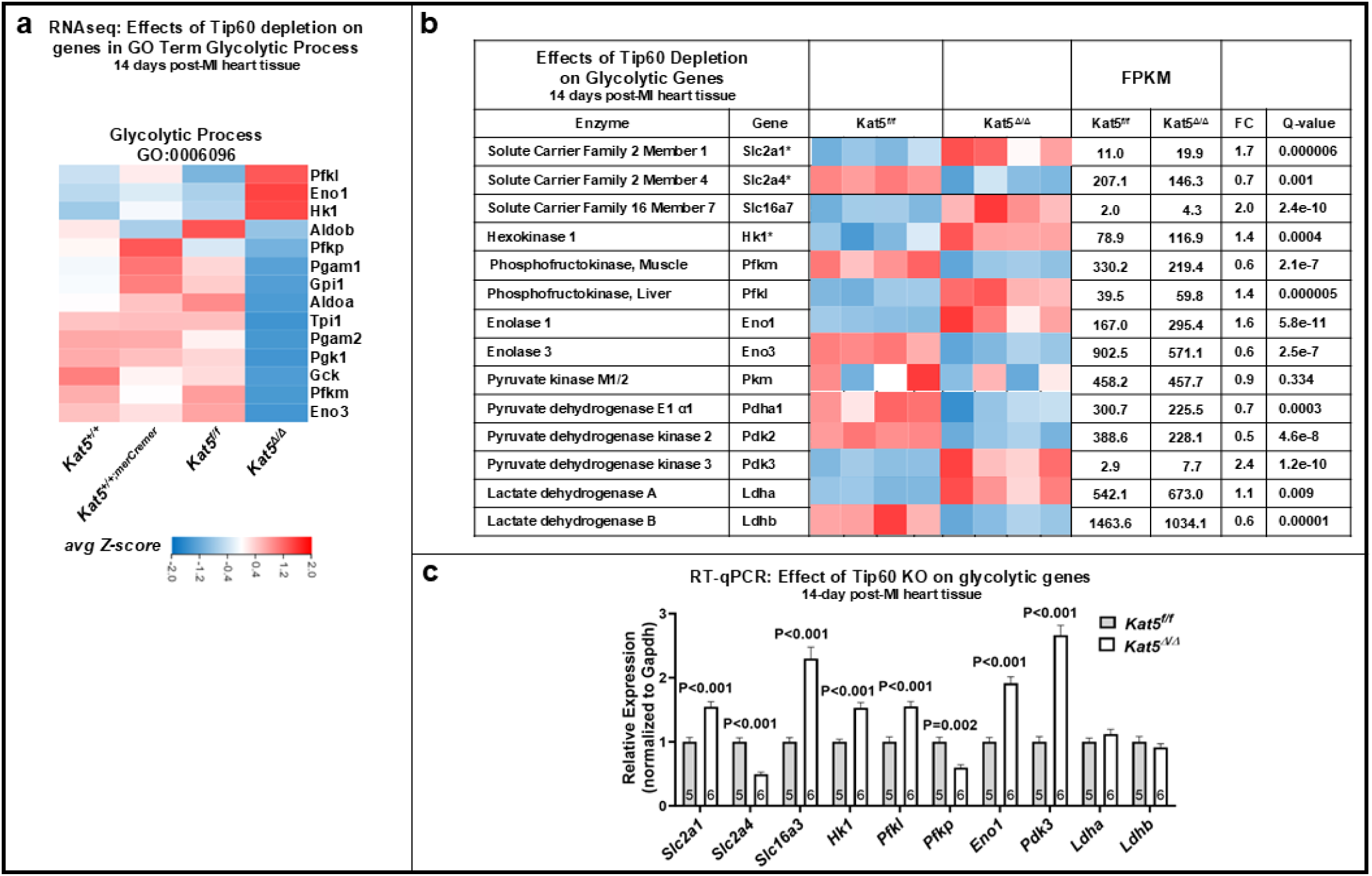
Expression of glycolytic genes in infarcted/Tip60-depleted hearts at 14 days post-MI. **Panel a**, RNAseq data showing mis-expressed genes in the GO term Glycolytic Process. *Kat5^f/f^*= control, *Kat5*^Δ*/*Δ^ = Tip60-depleted. *Kat5^+/+^*and *Kat5^+/+;merCremer^* assess off-target effects of Cre. **Panel b**, RNAseq data showing mis-expressed glycolytic genes at 14 days post-MI. Each column in the heatmap represents a biological replicate. **Panel c**, RT-qPCR verification of altered glycolytic gene expression in infarcted/Tip60-depleted hearts. Each biological replicate (N; shown inside bar) represents a single heart subjected to three technical replicates. Data are Mean ± SEM compared by unpaired, two-tailed Student’s t tests with Welch’s correction.

**Figure 8.**
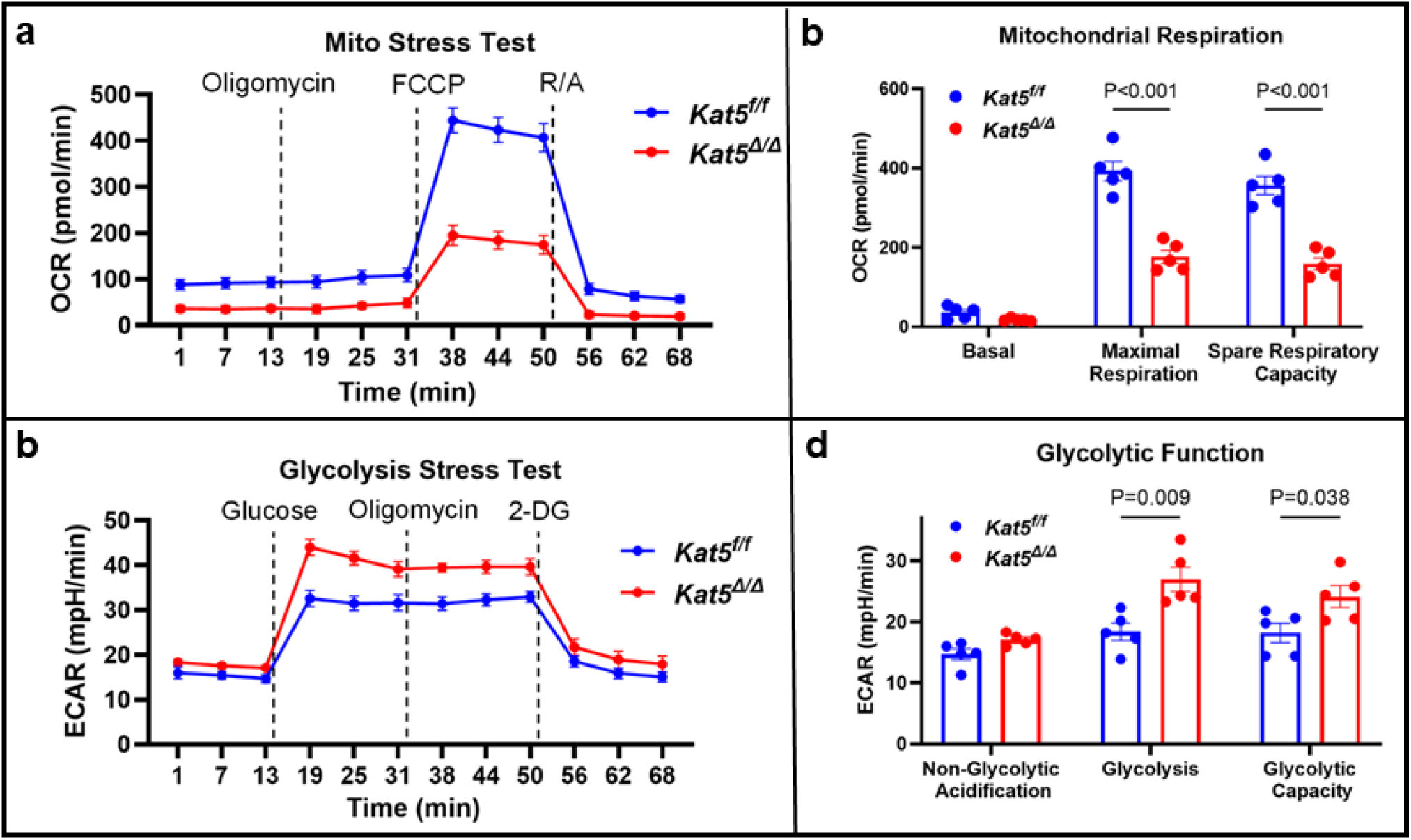
Seahorse assay indicating that Tip60 depletion induces metabolic reprogramming in isolated CMs characterized by reduced oxidative reserve and enhanced glycolysis. Extracellular flux analysis was performed with isolated CMs from naïve control (*Kat^5f/f^*) and experimental (*Kat5*^Δ*/*Δ^) mice. **Panels a-b:** Mito Stress Test (OCR, oxygen consumption rate; substrates, 1 mM pyruvate, 2 mM glutamine, 10 mM glucose). Representative OCR traces following sequential injection of oligomycin (ATP synthase inhibitor), FCCP (uncoupler), and rotenone/antimycin (Rot/A; Complex I/III inhibitors). **Panels c-d:** Glycolysis Stress Test (ECAR, extracellular acidification rate; substrate, 2 mM glutamine). ECAR traces following sequential injection of glucose, oligomycin, and 2-deoxyglucose (glycolysis inhibitor). OCR and ECAR parameters normalized to DNA content; Mean ±SEM compared using unpaired, two-tailed Student’s t tests with Welch’s correction; all Ns=5.

Regarding cell-cycle activation, post-MI depletion of Tip60 was associated with the up-regulation of individual genes in the Cell Cycle GO term (0007049; Fig. 2a), which was among the top 10 up-regulated GO terms overall (Supp. Fig. 2a). Regarding our recent report that depletion of Tip60 post-MI resulted in 2-3-fold in-creased expression of Sand G_2_-phase regulators *Ccna2*, *Ccnb1* and *Cdk1*^11^, RNAseq confirmed and extended these findings by revealing increased expression of the pro-proliferative genes *Top2a, Kif23, Mki67, Bub1, Prc1,* and *Stmn1* (Fig. 2b) in infarcted/Tip60-depleted hearts, all of which were recently shown to be upregulated in a proliferating subpopulation of CMs in the adult heart^34^.

### Altered Expression of Genes Indicative of EMT Transitioning & Cytoskeletal Disassembly in Infarcted/Tip60-depleted Hearts

Dedifferentiating CMs undergo a partial epithelial to mesenchymal – i.e. an EMT-like – transition^17,35^ accompanied by disassembly of the cytoskeleton/sarcomere complex^36^ (review^24^). As shown in Figure 3a, >75% of genes in the GO Term ‘Epithelial-Mesenchymal Transition’ were upregulated in infarcted/Tip60-depleted hearts, including 28 genes reported to be induced by over-expression of *Erbb2,* which promotes dedifferentiation and proliferation^17^ (Fig. 3b). RT-qPCR confirmed up-regulation of EMT-like genes, including *Actn1*^36^ and *Cttn,* which encodes cortactin to support cell migration^37–39^, as well as a trend toward increasing levels of *Nes* (nestin), which was recently shown to be increased in dedifferentiating CMs^17^ (Fig. 3c). Among cytoskeletal assembly genes, the decrease in *Mapt*, which encodes Tau to maintain microtubule stability and is selectively expressed in CMs^33^, is of interest due to its dependence on Tip60 during neurogenesis^29^. Other depressed cytoskeletal genes included *Mylk4* and *Myom1*, which maintain myofilaments^40,41^ ^14,21^. Regarding the observed depression of *Acta2* (Fig. 3b-c) we speculate that this reflects its down-regulation in non-CMs, since assessments in isolated CMs have shown that *Acta2* is upregulated in dedifferentiating CMs^14,16^, consistent with our immunostains indicating that α-SMA is increased within border zone CMs of infarcted/Tip60-depleted hearts (Supp. Fig. 7). Regarding the upregulated markers in Figure 3c-d it is noteworthy that *Cdk1* may regulate sarcomere disassembly^36^ and that *Shroom 3* promotes cytoskeletal organization^42^. Most importantly, and in accord with recent findings^43,44^, decreased expression of *Tnni3* in combination with the fourfold increase in *Tnni1* provides strong evidence that Tip60 KO induces a transition toward the immature dedifferentiated state (Fig.3c-d).

### Expression of ‘Soft’ ECM Genes in Infarcted/Tip60-depleted Hearts

As the heart develops, its extracellular matrix (ECM) transitions from relative softness toward rigidity. At immature stages, collagen 4, emilin-1, fibronectin and periostin are relatively abundant, followed by increased percentages of collagen-1, collagen-3 and laminin in adults^26^. In the neonatal heart, softness promotes ability of neonatal CMs to proliferate^18^. To facilitate CM proliferation in the adult heart, a metalloproteinase-induced transition toward soft ECM is required^45,46,47^. As shown in Supplemental Figure 8a, the expression of most genes in the GO Term ‘Extracellular Matrix’ was altered in Tip60 KO hearts, including increased levels of ‘softening’ genes in Supplemental Figure 8b-c (*Emilin1*, *Fn1*), among which *Cthrc1* and *Postn* have been respectively shown to promote ECM reconstruction^48,49^ and CM proliferation^50^. It is also notable that genes previously associated with Erbb2-induced CM dedifferentiation^35^ – *Fn1*, *Loxl2, Col3a1*, *Col4a2*, *Col6a1*, *Col8a2*, *Col15a1* – were increased (Supp. Fig. 8c), among which *Col15a1* has been shown to promote matrix remodeling and CM proliferation^51–53^. Tip60 KO also increased the expression of ECM genes associated with Fgfr1/Osmr-induced dedifferentiation^16^, including *Agrn* (agrin) which has been shown to promote regeneration post-MI^25,54^ (Supp. Fig. 8c).

### Altered Metabolic Programming in Infarcted/Tip60-depleted Hearts

During heart development, CM maturation is accompanied by a metabolic transition from glycolysis to FAO^55^. Post-MI, the ability of CMs to re-enter the cell-cycle requires reversion to glycolytic metabolism^16,17,19,56^. As shown in Figure 4, RNAseq revealed reduced expression of nearly all genes in the GO terms ‘Fatty Acid βOxidation’ (Fig. 4a). Downregulated FAO genes included those regulating the breakdown of short- thru very long-chain fatty acids (*Acads, Acadm, Acadl, Acadvl*; Fig. 4b-c), as well as reduced expression of carnityl palmitoyltransferase 1B (*Cpt1b*), KO of which was recently shown to be associated with dedifferentiation and renewed proliferation of adult CMs^19^.

Recent reports are relevant to these results. Most recently (2026), it was found that inhibition of Yap in the adult heart maintains oxidative metabolism in CMs by promoting fatty acid (FA) utilization, which is essential for maintaining cardiac maturation and function^57^. This finding was based, in part, by monitoring the expression of genes within a retinoic acid receptor (Rxr)-induced module including 16 “CM-intrinsic/RXR-controlled” genes that were previously shown to be regulated by embryonic Rxrs-α (Rxra) and -β (Rxrb), which are indispensable for vitality and maturation of CMs of the neonatal heart^58^. In a report addressing the role of PGC1/PPAR in CM maturation, it was shown that Rxra, and especially the ‘adult’ receptor Rxr-γ (Rxrg), were among the most strongly expressed transcriptional regulators in adult CMs^59^. In our study, nearly all genes within the “CM-intrinsic/RXR-controlled” module were repressed infarcted/Tip60-depleted adult hearts, as well as the expression of genes encoding Rxrg and the orphan receptor Rorc (Fig. 5a). In addition, Cut&Tag revealed that *Rxrg* and *Rorc* were among 47 CM genes (Fig. 10 below) exhibiting significant depletion of H2A.Zac^K4/K7^ in the promoter/TSS locus in Tip60-depleted hearts (Fig. 5b). To our knowledge, the role of Rxrg and Rorc in CM maturation has not been examined.

**Figure 9.**
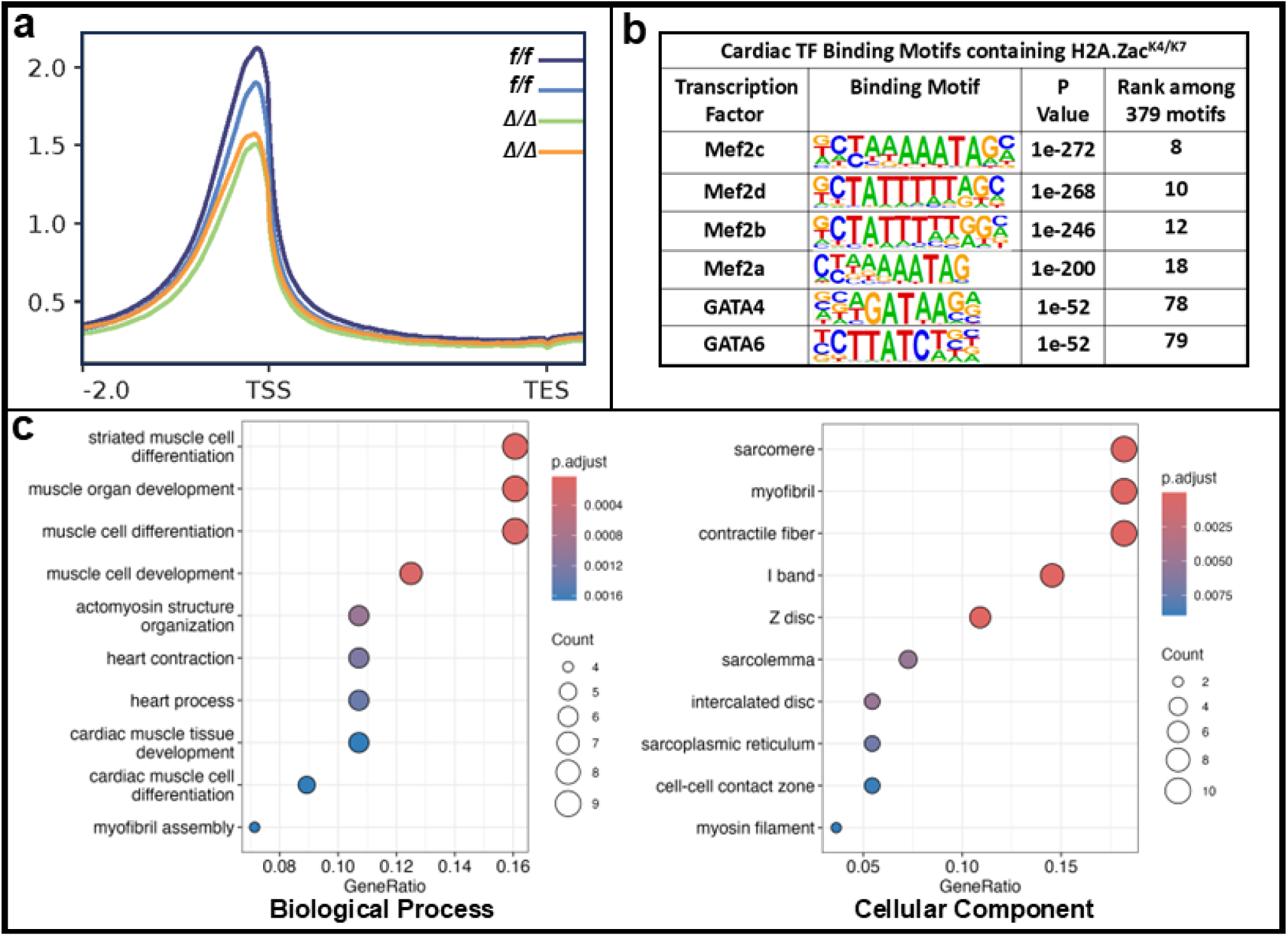
CUT&Tag performed on naïve/Tip60-depleted hearts reveals depleted levels of H2A.Zac^K4/K7^ in promoter/TSS loci of genes associated with CM maturity. **Panel a,** profile plots showing depletion of H2A.Zac^K4/K7^ in promoter/TSS loci. **Panel b**, HOMER analysis showing association of H2A.Zac^K4/K7^ with CM transcription factor motifs. **Panel c**, Gene Ontology (GO) dotplots indicating (at left) relatively high depletion of H2A.Zac^K4/K7^ in genes that regulate Biological Processes (BP) of striated muscle development and differentiation, and (at right) in Cellular Component (CC) genes associated with sarcomere maintenance. Results in panel **b** are based on p-value, in panels in **a** and **c** on adjusted p-value (p-adjust) to minimize false-positive when comparing results from only two biological replicates; *f/f*=control; Δ*/*Δ=Tip60 knockout in CMs.

**Figure 10.**
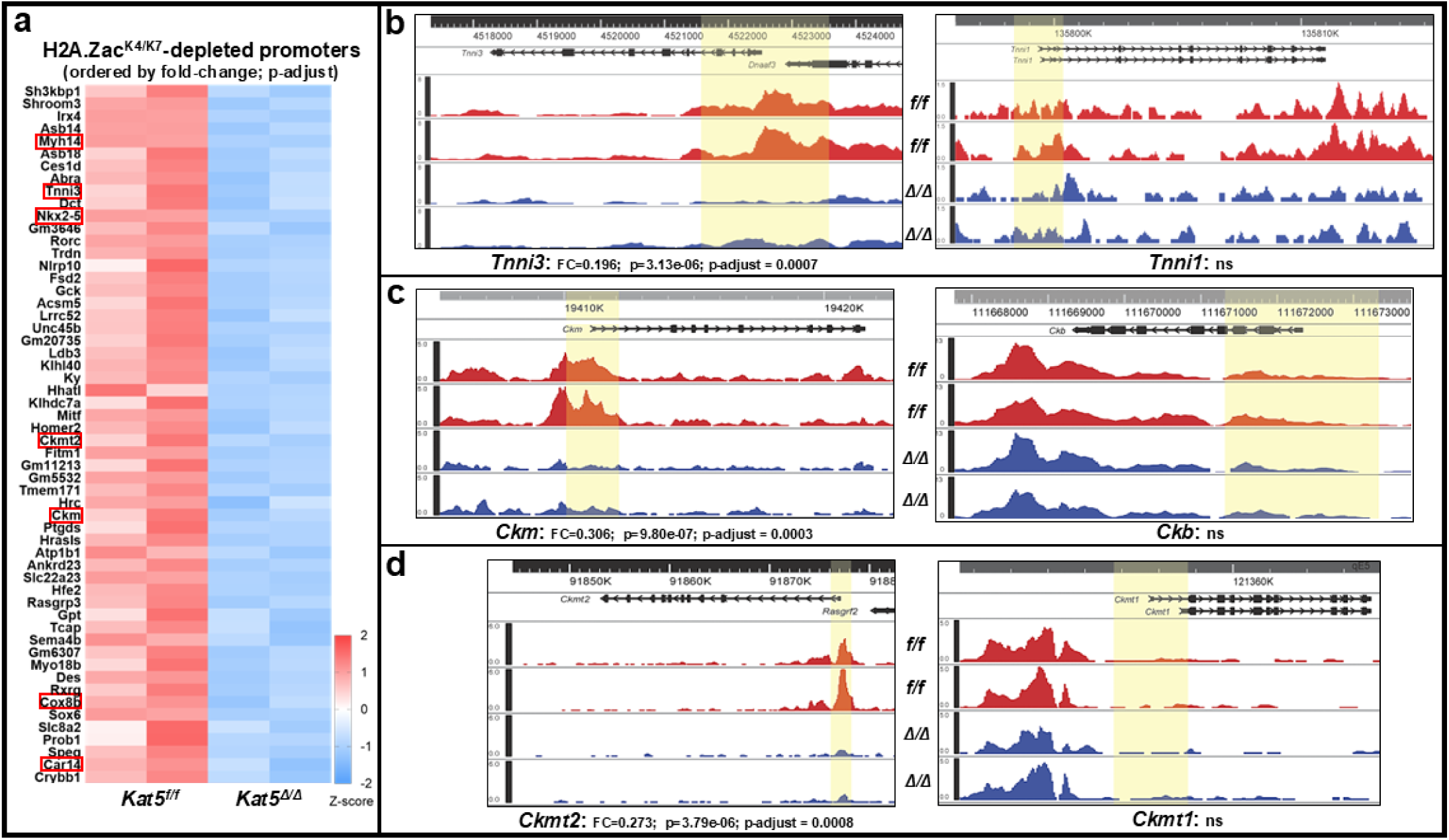
CUT&Tag: Effect of Tip60 depletion on H2A.Zac^K4/K7^ peaks in CM maturity genes. Tip60 was depleted in CMs of naïve adult mice and CUT&Tag was performed 11 days later. **Panel a**, heatmap showing 56 genes that exhibited statistically significant reduction, based on p-adjust values, of H2A.Zac^K4/K7^ in the promoter/TSS locus. **Panel b**, genome browser images showing reduced levels of H2A.Zac^K4/K7^ in the mature isoform of cardiac troponin I (*Tnni3*), but not in the embryonic isoform (*Tnni1*). **Panels c-d**: H2A.Zac^K4/K7^ peaks were also reduced in the adult (*Ckm; Ckmt2*), but not the embryonic (*Ckb; Ckmt1*), isoforms of creatine kinase (**c**) and mitochondrial creatine kinase (**d**). In **a**, red rectangles denote genes imaged by genome browsing in **bd** and in Supp. Fig. 9. For each genotype (control, *f/f*; Tip60 knockout, Δ*/*Δ), two biological replicates were performed, each consisting of nuclei isolated from the left ventricle, with statistical significance based on adjusted p-value (i.e., p-adjust) to minimize false-positives. *f/f*=control; Δ*/*Δ=Tip60 knockout. Yellow highlighting in **b-d** denotes ±1,000 bp around the TSS of each gene.

Regarding Krebs Cycle genes (Fig. 6a), KO of Tip60 strongly reduced the expression of nearly all isoforms within families that regulate each step of the cycle (Fig. 6b-c), among which inhibition of succinate dehydrogenase (*Sdh*) has been shown to promote CM proliferation and revascularization^60^. A notable exception to reduced expression of TCA cycle genes was the absence of an effect on α*-ketoglutarate dehydrogenase* (*Ogdh*), because its targeted disruption increases cell-cycle activation and numbers of CMs^19,61^.

Because genes regulating fatty acid β-oxidation (Figs. 4,5) and the TCA Cycle (Fig. 6) were reduced by Tip60 KO, it was anticipated that reversion to glycolysis would be observed. However, most genes in the GO term ‘Glycolytic Process’ were decreased (Fig. 7a), although those regulating crucial glycolytic steps (*Hk1*; hexokinase-1) as well as isoforms in glycolytic gene families that are normally modestly expressed – *Slc2a1*, *Slc16a7*, *Pfkl*, *Eno1*, *Pdk3*, and *Ldha* – were upregulated (Fig. 7b-c). To directly assess whether depletion of Tip60 promoted reversion toward glycolytic metabolism, Seahorse Stress Testing was performed. As shown in Figure 8a-b, while basal oxygen consumption rates (OCR) in CMs isolated from Tip60-depleted hearts were not significantly reduced, reduction of Maximal Respiration and Spare Respiratory Capacity was significant and accompanied by significant increases in glucose-mediated acidification of the extracellular environment (ECAR) and glycolytic capacity (Fig. 8c-d). These changes are consistent with a reversion toward glycolytic metabolism. In summary, data in Figures 4–8 support the hypothesis that depletion of Tip60 promotes an oxidative glycolytic transition – the latter possibly mediated via up-regulation minor isoforms – that is required for CM proliferation.

### Effects of Tip60 Depletion on H2A.Zac^K4/K7^ Levels in Cardiomyocyte Maturity Genes

To assess effects of Tip60 depletion on H2A.Zac^K4/K7^ levels in promoter/TSS loci, CUT&Tag was performed on naïve adult hearts. As shown in Figure 9, Tip60 KO diminished peaks in the promoter/TSS loci (Fig. 9a) of genes enriched in cardiac transcription factor binding sites (Fig. 9b). Remarkably, Gene Ontology (GO) analysis revealed that loss of H2A.Zac^K4/K7^ in promoter/TSS loci mostly occurred in genes that regulate sarcomere maintenance and muscle differentiation (Fig. 9c). Among 1,724 H2A.Zac^K4/K7^ peaks that were significantly diminished (per P-adjust) genome-wide, only 56 were localized in the promoter/TSS locus of individual genes (Fig. 10a), among which 47 were judged as ‘CM genes’ (i.e., cardiac-enriched; Supp. Table 3) based on their relative expression compared with all cell types present in heart tissue as listed in the Tabula Muris database^33^. Correlation of H2A.Zac^K4/K7^ depletion in CM genes with effects on their transcription in infarcted/Tip60-depleted hearts indicated that 48% exhibited reduced expression, favorably comparing with the effect of H3K4^me3^ reduction in *Cpt1b*-depleted CMs^19^, while six genes (including *Shroom3*, *Tcap*, & *Desmin*) were up-regulated. It was also of interest that *Mhrt*, which encodes a long non-coding RNA that limits expression of *Myh7* and cardiac remodeling^62^, was one of only two genes that exhibited increased levels of H2A.Zac^K4/K7^ (Supp. Table 3).

Among the 47 CM genes shown to exhibit reduced H2A.Zac^K4/K7^ (Fig. 10a), *Tnni3, Ckm, Ckmt2, and Cox8b* represent the mature isoform within their respective gene families. Comparison of genome browser images of these isoforms with those of their respective embryonic precursors (*Tnni1*, *Ckb*, *Ckmt1*, *Cox8a*) showed that acetylated H2A.Z was not diminished in any of the latter (Fig. 10b-d, Supp. Fig. 9a). It is noteworthy that cardiac myosin light chain (*Myl3*), expression of which is indicative of ventricular maturity, exhibited reduced levels of H2A.Zac^K4/K7^ in promoter/TSS loci of Tip60-depleted hearts, in contradistinction to levels in the *Myl1* isoform (Supp. Fig. 9a). Also, among several ‘gene model’ (Gm) loci exhibiting reduced H2A.Zac^K4/K7^, Gm6307, which encodes the lncRNA *Mir1a-1hg* (Supp. Fig. 10), is of interest because it harbors the bicistronic construct encoding microRNAs *Mir1a-1* and *Mir133a-2*, the latter having been shown to inhibit CM proliferation in the developing heart^63^; their role in heart development, function and pathology has been reviewed^64^.

Regarding effects of H2A.Zac^K4/K7^ depletion on gene expression, all mature isoforms were downregulated in infarcted/Tip60-depleted hearts (Supp. Table 3). Regarding expression of the embryonic isoforms, *Ckb* and *Myl1* were not affected, *Cox8a* was downregulated, and *Tnni1* (Fig. 3c-d) and *Ckmt1* were strongly upregulated (Supp. Tables 3-4). Taken together, results in Figures 5 and 9–10 are consistent with the possibility that an intact Tip60 H2A.Zac^K4/K7^ axis is necessary to maintain the expression of genes that confer cardiac maturity. In this regard it is noteworthy that Tip60 knockout in CMs did not significantly affect H2A.Zac^K4/K7^ levels in any genes that encode cyclins or cyclin-dependent kinases; examples are shown in Supplemental Figure 11.

## DISCUSSION

*Resume*: Our recent findings support the hypothesis that genetic depletion^10,11^ or pharmaceutical inhibition^12,13^ of the pleiotropic acetyltransferase Tip60 (*Kat5*) after myocardial infarction (MI) confers cardioprotection and regeneration by preventing the site-specific acetylation and activation of its targets, which are pro-apoptotic (i.e., p53) and anti-proliferative (i.e., Atm). Here we extend these findings by showing that post-MI depletion of Tip60 essentially eliminates site-specific acetylation of the histone variant H2A.Z (i.e. H2A.Zac^K4/K7^) in CMs, a chromatin target associated with induction and maintenance of the differentiated state in other cell-types. Western blotting indicated that depletion of Tip60 post-MI reduced levels of H2A.Zac^K4/K7^ in cardiac tissue by approximately two-thirds (Fig. 1b), and, most remarkably, immunostaining revealed near-complete loss of H2A.Zac^K4/K7^ in CM nuclei (Fig. 1a).

Elimination of H2A.Zac^K4/K7^ was associated with dedifferentiation (Figs. 2–7), a process now considered as prerequisite to the ability of CMs to undergo cell-cycle reactivation and proliferation to effect myocardial regeneration^14–18,24,65–67^. This result is consistent with recent findings that Tip60-specific acetylation of H2A.Z is required to maintain the differentiated state of HSCs^28^, and neurons^29^; because these reports showed that acetylated H2A.Z binds pioneer transcription factors that activate genes to establish cellular maturity – specifically Myc in HSCs^28^ and Ascl1 in neurons^29^ – it will be of interest to assess whether H2A.Zac^K4/K7^ recruits cardiomyogenic transcription factors, consistent with HOMER results (Fig. 9b; Supp. Fig. 1c), to enhancer/promoter loci of genes that maintain the CM differentiated state. Speculation that acetylation of H2A.Z is required to specify *all* cell-types^68^ is consistent with earlier findings showing that H2A.Z acetylation is required for skeletal^69^ as well as smooth^70^ muscle cell differentiation, as well as the recently published finding that Tip60 drives cardiac fibroblast myofibroblast differentiation, and potentially scar formation^71^. A crucial role for Tip60 within selective genomic loci as cells differentiate is consistent with recently published cryo-electron microscopic findings that disruption of the NuA4/Tip60 complex results in genome-wide dispersion of acetylated H2A.Z^72^.

### Extracellular Matrix Transition

As shown in Supplemental Figure 8, ECM genes associated with the transition to soft neonatal-like matrix, which promotes CM migration and proliferation^16–18^, exhibited increased expression after Tip60 depletion. While it is unclear why genes associated with stiffening and scarring – *Col1a1*, *Col3a1*, *Lox*, *Loxl2* – were also upregulated, *Col1a1* and *Lox* have been associated with CM dedifferentiation^16^. It is also interesting to consider that matrix softening is enhanced by the large increase in *Cthrc1* (collagen triple helix repeat containing-1), a secreted glycoprotein that has been shown to mediate wound healing^49^, limit matrix deposition during arterial remodeling^73,74^, and promote cell migration^75^. An unresolved question regards why, when Tip60 is specifically depleted in CMs, ECM genes that are selectively expressed in fibroblast subpopulations^49^ transition toward softness. Although transition toward soft ECM has been shown to be regulated by crosstalk between the Wnt/β-catenin, Hippo/Yap and TGFβ pathways^76^, mechanistic details regarding signaling between the cells involved remains speculative.

### Metabolic Reprogramming

Effects of depleting Tip60 on the expression of metabolic genes (Figs. 4–8) are consistent with multiple findings that CM dedifferentiation is accompanied by metabolic reprogramming, from FAO toward glycolysis (review^77^). Most prominently, Tip60 depletion was associated with reduced expression of genes regulating key steps in the TCA cycle (Fig. 6) and FAO (Figs. 4,5). Recent findings strongly show that FAO metabolism is required to attain and maintain the differentiated state of CMs, reversion from which is required to permit their proliferation and potential regeneration^19,57,58^. In this regard it will be of interest to assess whether the retinoic acid receptor *Rxrg* and the orphan receptor *Rorc*, which induces differentiation in Th17 cells^78,79^, promote the differentiated state in CMs by inducing the expression of CM-intrinsic RXR-controlled FAO genes.

While RNAseq evidence for renewed glycolysis in Tip60-depleted hearts was not definitive (Fig. 7a), previous observations that reversion to glycolysis during CM dedifferentiation is accompanied by increased expression of genes encoding pyruvate dehydrogenase kinase 3 (*Pdk3*), monocarboxylate transporter 3 (*Slc16a3;* Mct4)^17^, and glucose transporter 1 (*Slc2a1*; Glut1)^56^ were recapitulated (Fig. 7b-c). Although the most robustly expressed glucose transporter in CMs, *Slc2a4* (Glut4), was depressed by Tip60 depletion, expression of the weakly expressed isoform *Slc2a1* was upregulated (Fig. 7b-c), which may be relevant since over-expression of *Slc2a1* has been shown to mitigate heart failure^80^ and more recently associated with cardiac regeneration^81,82^. Post-MI depletion of Tip60 was also associated with significant up-regulation of isoforms in glycolytic gene families (*Slc16a7*, *Pfkl*, *Eno1*, *Eno2, Ldha;* Fig. 7b-c) that are normally weakly expressed in the adult heart. Taken together, these findings suggest that reversion toward glycolysis is mediated by the increased expression of isoforms that are normally weakly expressed in the adult heart, a possibility consistent with Seahorse results (Fig. 8).

### Effect of Tip60 Depletion on CM Proliferation

Our previous findings that Tip60 depletion post-MI maintains cardiac function, diminishes fibrosis and activates the CM cell-cycle^10,11^, as well as the findings reported here that depletion induces a dedifferentiated state permissive for CM proliferation, are consistent with the notion that targeting Tip60 permits heart regeneration post-MI. However, it remains unsettled whether bona fide proliferation of CMs ensues. Per our recent review^83^, resolution of this seemingly straightforward question has been forestalled by inadequate methodology to definitively ensure that CMs exhibiting cell-cycle markers in the *in vivo* heart are (i) indeed CMs that (ii) are not polyploid. We are rigorously addressing this question using updated methodology designed to provide unequivocal assessment of CM identity, ploidy and nuclearity^84^.

### Epigenetic Mechanisms to Maintain the Differentiated State

It was recently reported that accumulation of αketoglutarate (αKG) resultant from CM-specific depletion of *Cpt1b* activates lysine demethylase 5 (Kdm5), thereby reducing histone H3K4^me3^ in CM maturity genes to levels that cause CM dedifferentiation and cellcycle activation^19^. Subsequent findings indicate that systemic administration of αKG post-MI has similar effects^61^. Although depletion of Tip60 did not affect the expression of *Idh2* and *Ogdh* in a fashion predicted to increase αKG levels (Fig. 6b,c), immunostaining revealed modest reduction of H3K4^me3^ levels in CM nuclei (Supp. Fig. 12). Findings that Tip60-acetylated H2A.Zac^K4/K7^ in combination with H3K4^me3^ comprise a binary signature that maintains expression of neuronal maturity genes^68^ are consistent with our findings, which are being extended to assess the effects of depleting Tip60 on this binary signature, as well as on alternatively modified histone H3 in promoter/TSS loci of CM maturity genes.

Results in Figures 9–10 are consistent with the possibility that a Tip60 H2A.Zac^K4/K7^ axis maintains the differentiated state in CMs. Tip60 knockout strongly increased expression of the immature isoform *Tnni1* (Fig. 3c-d), while reducing levels of H2A.Zac^K4/K7^ in the promoter /TSS, and inhibiting expression, of the adult Tnni3 isoform (Fig. 3c-d). This is perhaps of most relevance since *Tnni3* has been considered as a cardinal indicator of CM maturation^43,44^. Tip60 KO similarly affected H2A.Zac^K4/K7^ levels in the mature isoforms of other CM genes that possess embryonic isoforms, including *Ckm*, *Ckmt2*, and *Cox8b* (Fig. 10c-d; Supp. Fig 9a). However, depletion of H2A.Zac^K4/K7^ in combination with reduced expression was not observed for other genes that have been considered markers of CM maturity. For example, as shown in Supplemental Table 3, among 12 H2A.Zac^K4/K7^depleted genes considered to be 90-100% CM-specific by Tabula Muris, only 9 were down-regulated in infarcted/Tip60 depleted hearts. Among these, although KO reduced H2A.Zac^K4/K7^ in the genes encoding cardiac transcription factors Nkx2-5 and Ppargc1a^85^ (which notably heterodimerizes with Rxrs), their expression was respectively increased and decreased (Supp. Table 4). Considering the complexities of epigenetic regulation, which include modifications to histone H3 that either activate (H3^K4me3^, H3^K27ac^) or silence (H3^K27me3^) transcription, as well as possible effects of p300 (*Kat3b*) which also acetylates lysines 4 and 7 in H2A.Z^85,86^, this was not unexpected. Also, negative feedback signaling promoting CM redifferentiation^15,17,87,88^ may have contributed to the non-uniform response of CM maturity genes. For example, increased expression of all transiently upregulated angiogenic inhibitors, plus several tumor suppressors (*Phlda1*, *Phlda3*, *Numbl*, *Nab2*) shown to accompany redifferentiation^17^, was observed in Tip60-depleted hearts at day 14 post-MI. Although hippo pathway components were unaffected (Supp. Table 5), transient up-regulation may have occurred prior to the 14-day post-MI timepoint employed in our model, a possibility that is being followed-up. We conjecture that negative feedback signaling also explains the absence of an effect on *Ogdh* expression (Fig. 5), as well as why the expression of cardiac transcription factor genes (*Gata4*, *Hand2*, *Myocd*, *Nkx2-5)* was upregulated in infarcted/Tip60-depleted hearts, especially since H2A.Zac^K4/K7^ was significantly depleted in *Myocd* and *Nkx2-5* promoters (Supp. Table 4).

In summary, considered within the context of findings in other cell types^28,68–70,72,89–93^ – and in particular the very recent and highly relevant demonstration that Tip60 H2A.Zac axis may be a major driver of myofibroblast-mediated fibrosis^71^ – our results support the hypothesis that site-specific acetylation of H2A.Z by Tip60 maintains the expression of genes that drive and maintain the differentiated state in CMs. Considered along with our previous findings that depletion of Tip60 in CMs beginning three days post-MI promotes cell-cycle activation, reduces apoptosis and scarring, and preserves cardiac function^10,11^ – effects that are mimicked using drugs that inhibit Tip60’s acetyltransferase (AT) domain^12,13^ – pharmaceutical targeting of this pleiotropic factor is warranted to ameliorate cardiac injury.

## Non-Standard Abbreviations

Cre: Cre-recombinase;
CM: cardiomyocyte;
cTnT: cardiac troponin-T;
DEG: differentially expressed gene;
FAO: fatty acid oxidation;
H2A.Zac^K4/K7^: histone H2A.Z acetylated at lysines 4 and 7;
KO: *Kat5* (lysine acetyltransferase-5; Tip60) genetic knockout;
MI: myocardial infarction;
Tip60: Tatinteractive protein 60 kD protein.

## SUPPLEMENTAL FIGURES & LEGENDS

**Supplemental Figure 1.**
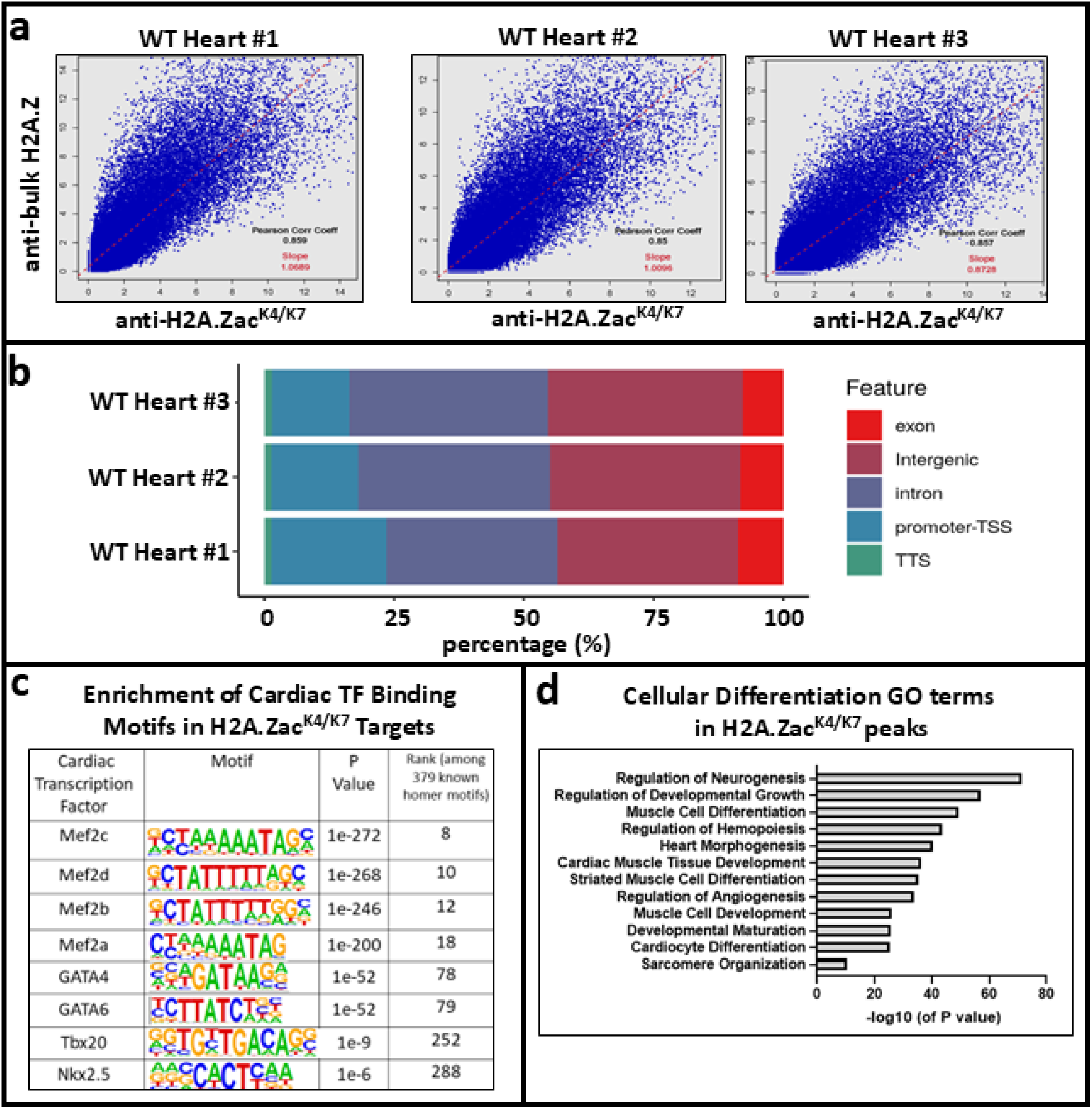
CUT&Tag genomic localization of H2A.Zac^K4/K7^. Panel. **a**: Qualification of antiH2A.Zac^K4/K7^ antibody (Cell Signaling Technology #75336) for ability to perform CUT&Tag on adult heart tissue by comparing targets recognized by commercially qualified antibody anti-bulk H2A.Z (Active Motif #39013). **Panel b**: Percentage of H2A.Zac^K4/K7^ peaks in annotated genomic loci. **Panel c**: Cardiac TF binding motifs enriched in H2A.Zac^K4/K7^; identified using Homer. **Panel d**: Gene Ontology (GO) terms associating muscle cell differentiation and development with H2A.Zac^K4/K7^ peaks. CUT&Tag was performed on triplicate Wild-Type hearts that were not infarcted.

**Supplemental Figure 2.**
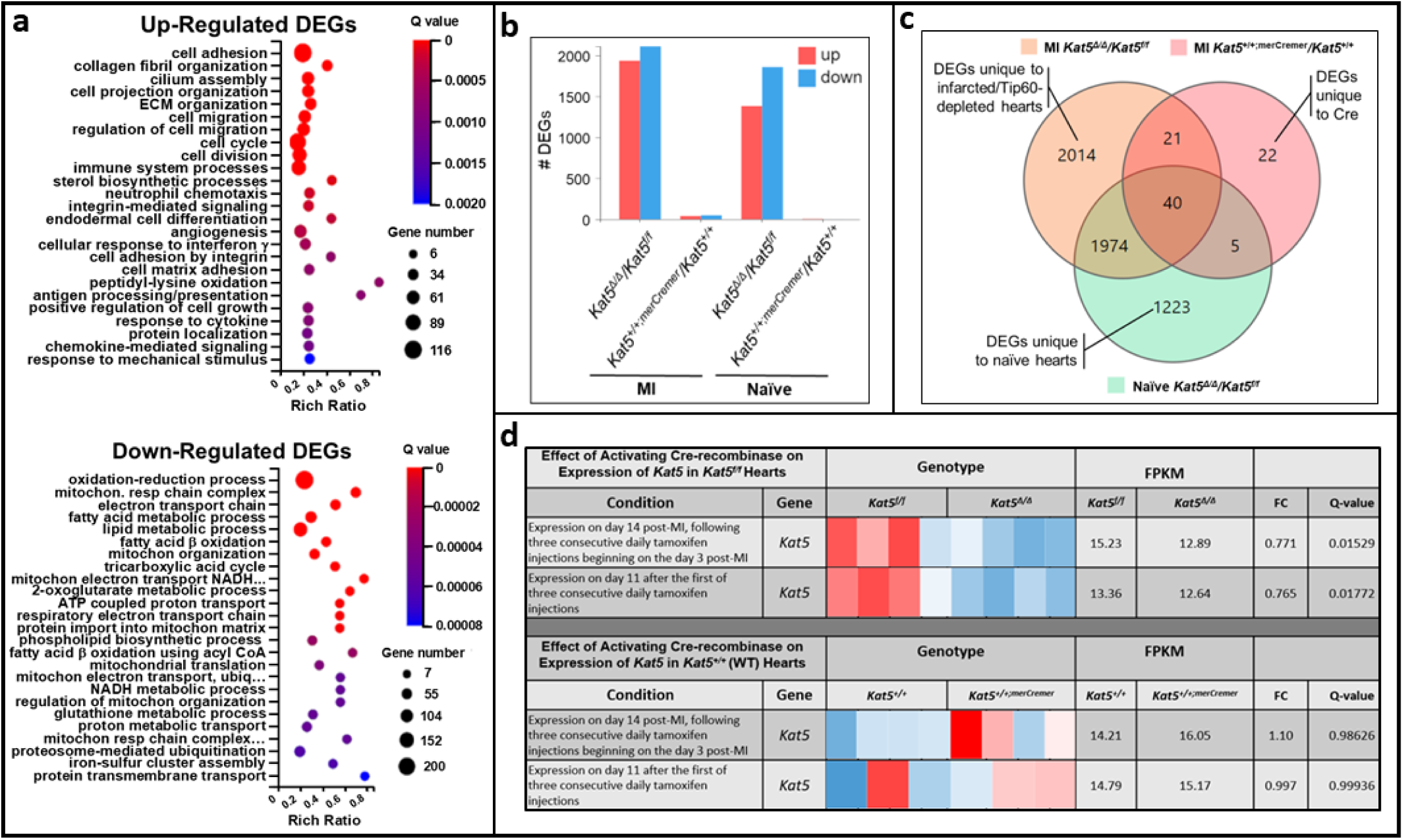
RNAseq data showing transcriptional effects of Tip60 depletion, and of Crerecombinase side-effects in the absence of gene ablation. Twelve-week-old *Kat5^f/f^*v. *Kat5*^Δ^*^/^*^Δ^ mice or *Kat5^+/+^* v. *Kat5^+/+;merCremer^* mice were intraperitoneally injected with 40 mg/kg tamoxifen for three consecutive days. In experiments performed on infarcted mice, tamoxifen was administered for three consecutive days beginning on the third day post-MI, followed by collection of hearts on the 14^th^ day post-MI. Naïve mice of similar age were identically treated, followed by collection of hearts 11 days after the first injection. **Panel a**: Top 25 up- and down-regulated GO terms in infarcted/Tip60-depleted hearts. Rich Ratio denotes DEGs/total genes; bubble size denotes # DEGs annotated to GO term; colors denote statistical significance. **Panel b:** Number of up/downregulated DEGs in infarcted (MI) and naïve hearts, in the presence (*Kat5*^Δ^*^/^*^Δ^) and absence (*Kat5^+/+^*) of Tip60 knockout; notably, off-target effects of Cre were minimal. **Panel c**: Venn diagram showing DEG overlap between groups. **Panel d**: RNAseq data showing that expression of the *Kat5* gene is unaffected when Crerecombinase is activated in wild-type (*Kat5^+/+^*) mice; each colored column represents a biological replicate.

**Supplemental Figure 3.**
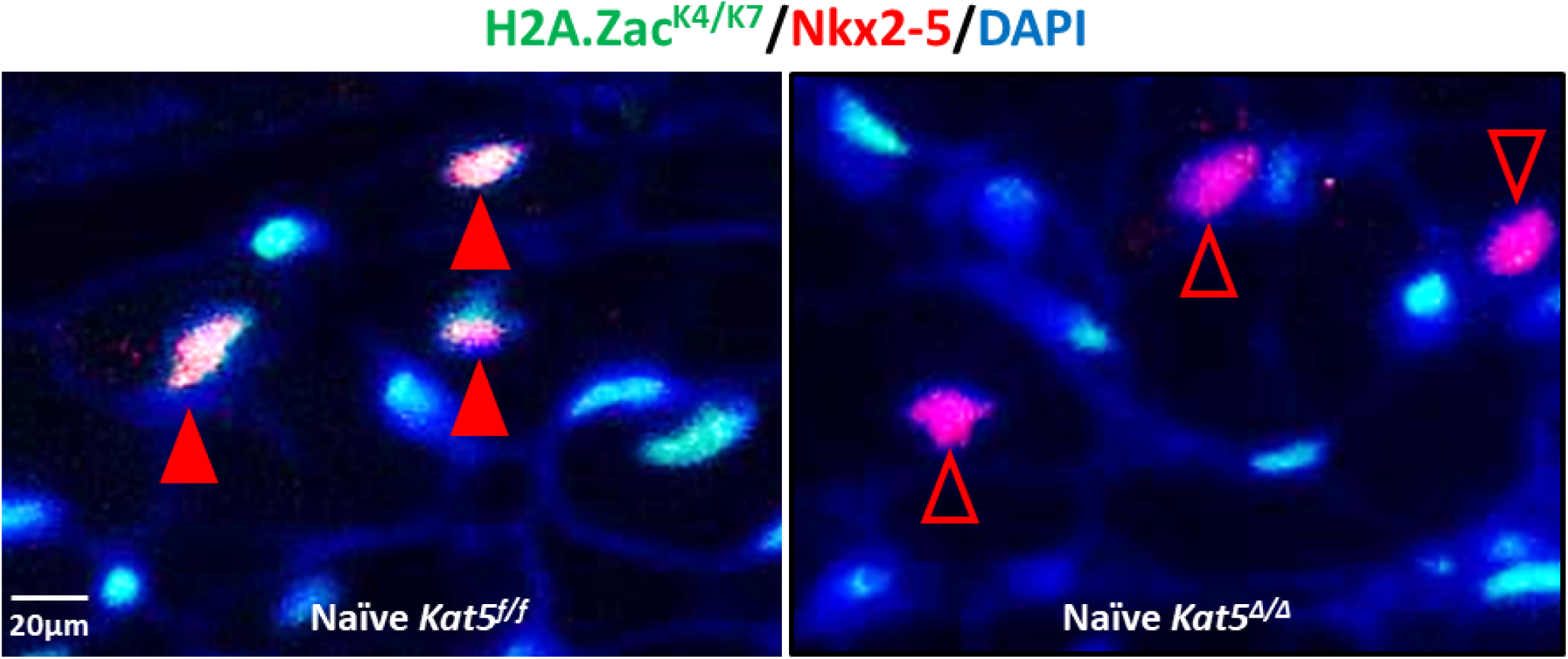
Verification of cardiomyocyte identity by immunostaining Nkx2-5. Left: Closed red arrowheads in control (*Kat5^f/f^*) myocardium denote CM nuclei immuno-stained for Nkx2-5 (red fluorescence) that contain site-specifically acetylated H2A.Z (H2A.Zac^K4/K7^; green fluorescence). **Right:** Open red arrowheads denote Nkx2-5-positive CM nuclei devoid of H2A.Zac^K4/K7^ when Tip60 is depleted (*Kat5*^Δ^*^/^*^Δ^).

**Supplemental Figure 4.**
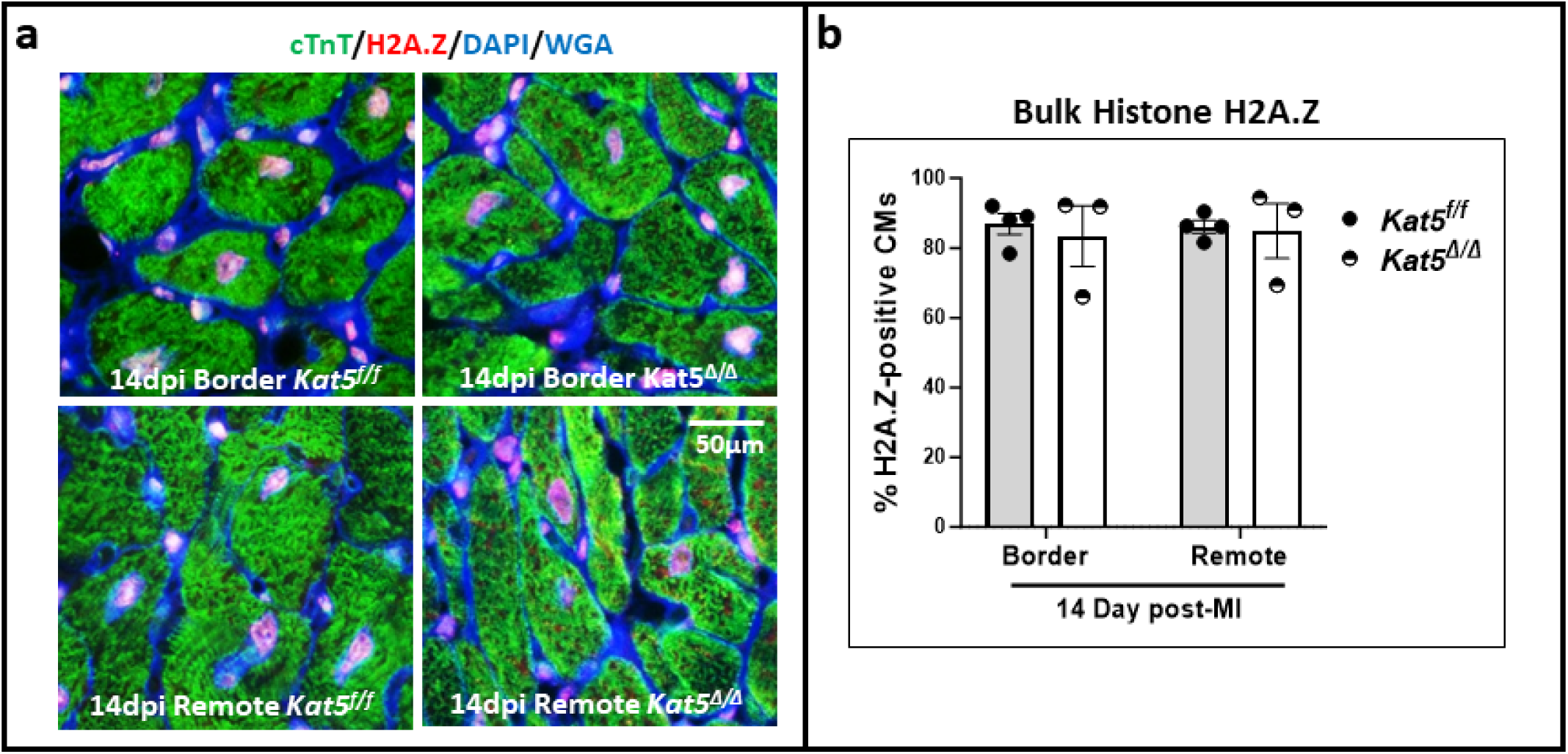
Percentages of CMs containing bulk histone H2A.Z are unaffected by Tip60 depletion. Panel. **a**, immunostaining of bulk histone H2A.Z (red fluorescence) in CM nuclei; CMs are identified by cardiac troponin-T immuno-staining. **Panel b**, percentage of CMs containing bulk H2A.Z. In each biological replicate, six cross-sectional fields are assessed in the border and remote zone at 400x magnification. P<0.05 vs. *Kat5^f/f^* using unpaired Student’s t-test; means ±SEM.

**Supplemental Figure 5.**
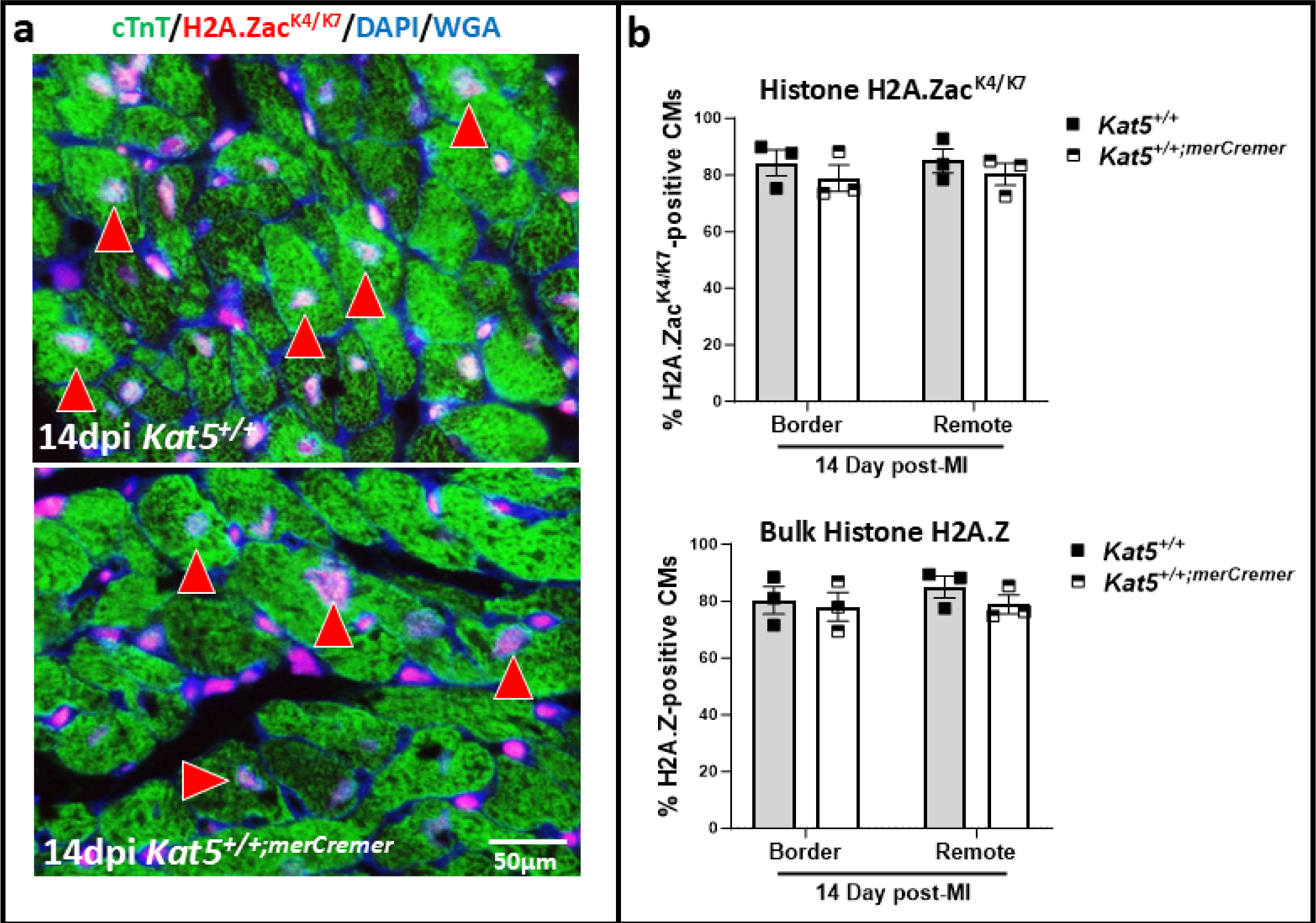
Cre does not affect numbers of CMs that contain H2A.Zac^K4/K7^ or bulk histone H2A.Z in infarcted hearts. Panel a, above: Wild-type CMs not expressing Cre (*Kat5^+/+^*); immunostaining shows H2A.Zac^K4/K7^-positive (red fluorescence) nuclei in wild-type CMs. **Panel a, below**: Wild-type CMs expressing Cre (*Kat5^+/+;merCremer^*). In both panels red arrowheads denote H2A.Zac^K4/K7^-positive CMs. **Panel b** shows enumeration of H2A.Zac^K4/K7^-positive CMs (**above**), and bulk H2A.Z-positive CMs (**below**). CMs are identified by anti-cardiac troponin-T (green fluorescence) immunostaining. P<0.05 vs. *Kat5^+/+^*per unpaired Student’s t-test; means ±SEM.

**Supplemental Figure 6.**
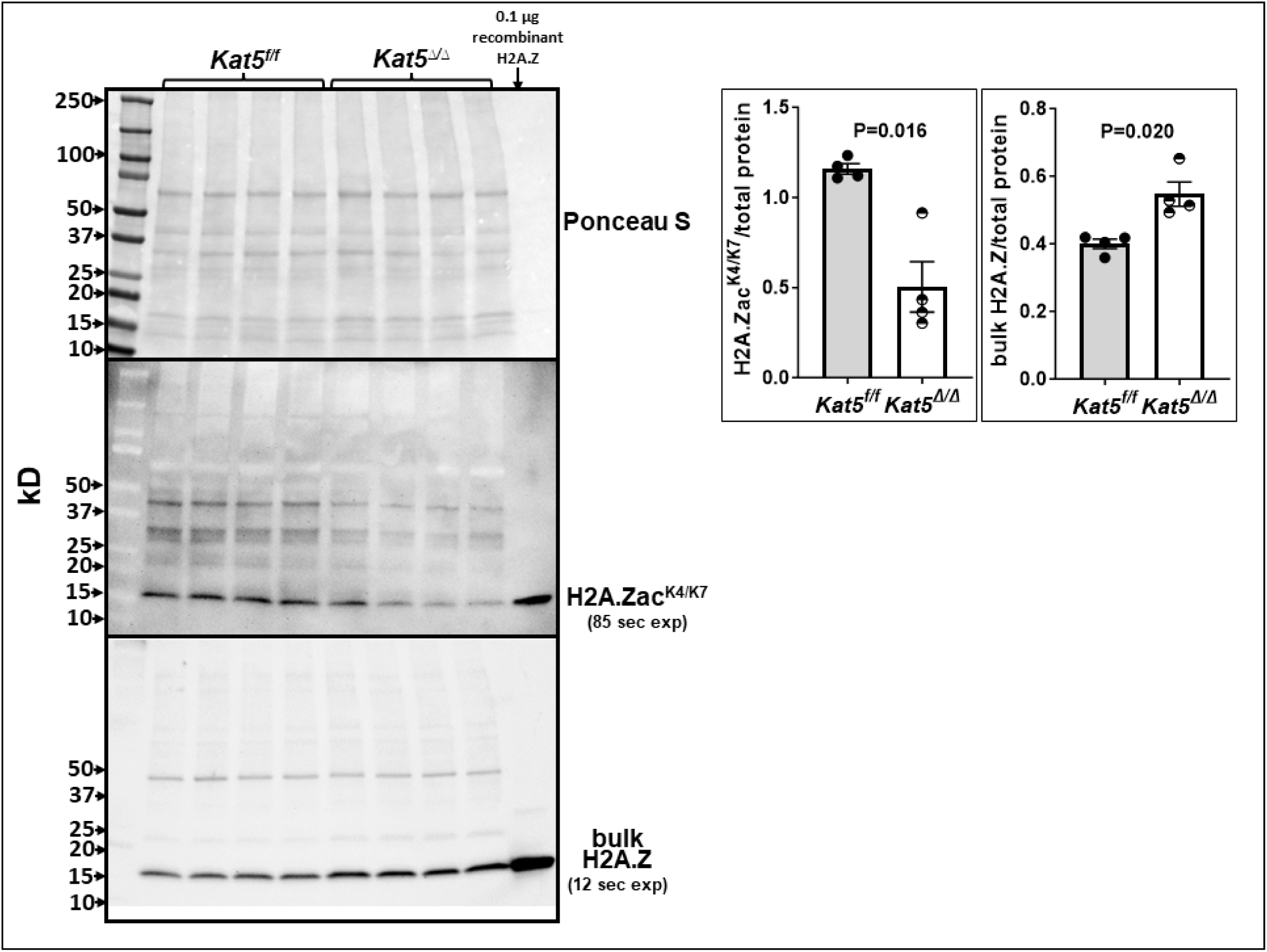
Non-cropped western blots showing reduced levels of H2A.Zac^K4/K7^ in Tip60depleted (*Kat5*^Δ^*^/^*^Δ^) hearts. In the bar graphs (right), each point represents a biological replicate (i.e., an individual mouse heart).

**Supplemental Figure 7.**
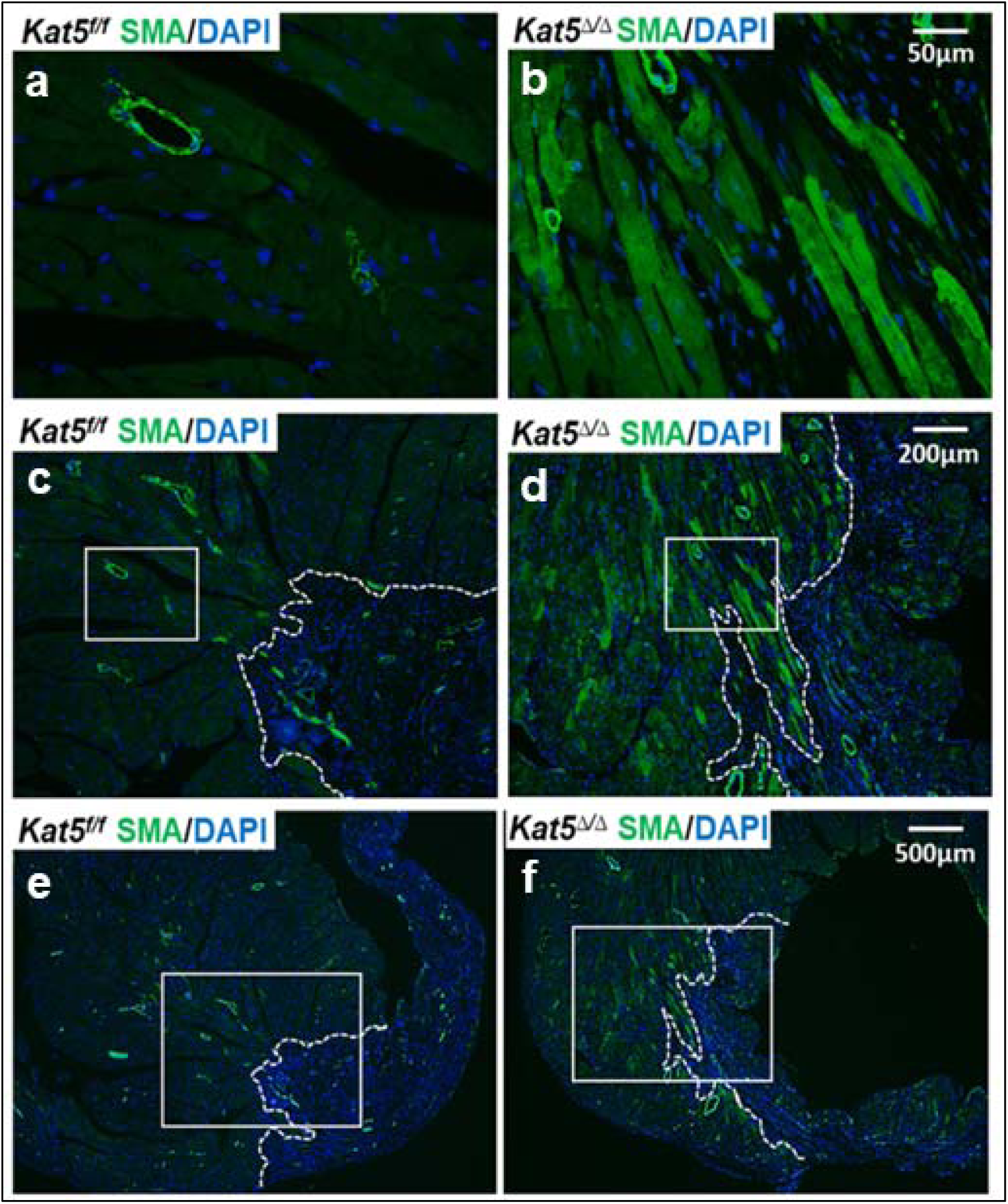
α-SMA-positive CMs in the border zone of infarcted/Tip60-depleted hearts. **Panels a-f** are representative α-SMA-immunostained images of control (Kat5^f/f^; left column) and Tip60depleted (Kat5^Δ/Δ^; right column) hearts at 28 days post-MI. Striated α-SMA-positive CMs are aligned along the border zone of Tip60-depleted hearts. **Panels a-b** are shown at reduced magnification in the white rectangles in panels c-f. The broken white line in panels c-f demarcates the border (left) and infarct (right) zones.

**Supplemental Figure 8.**
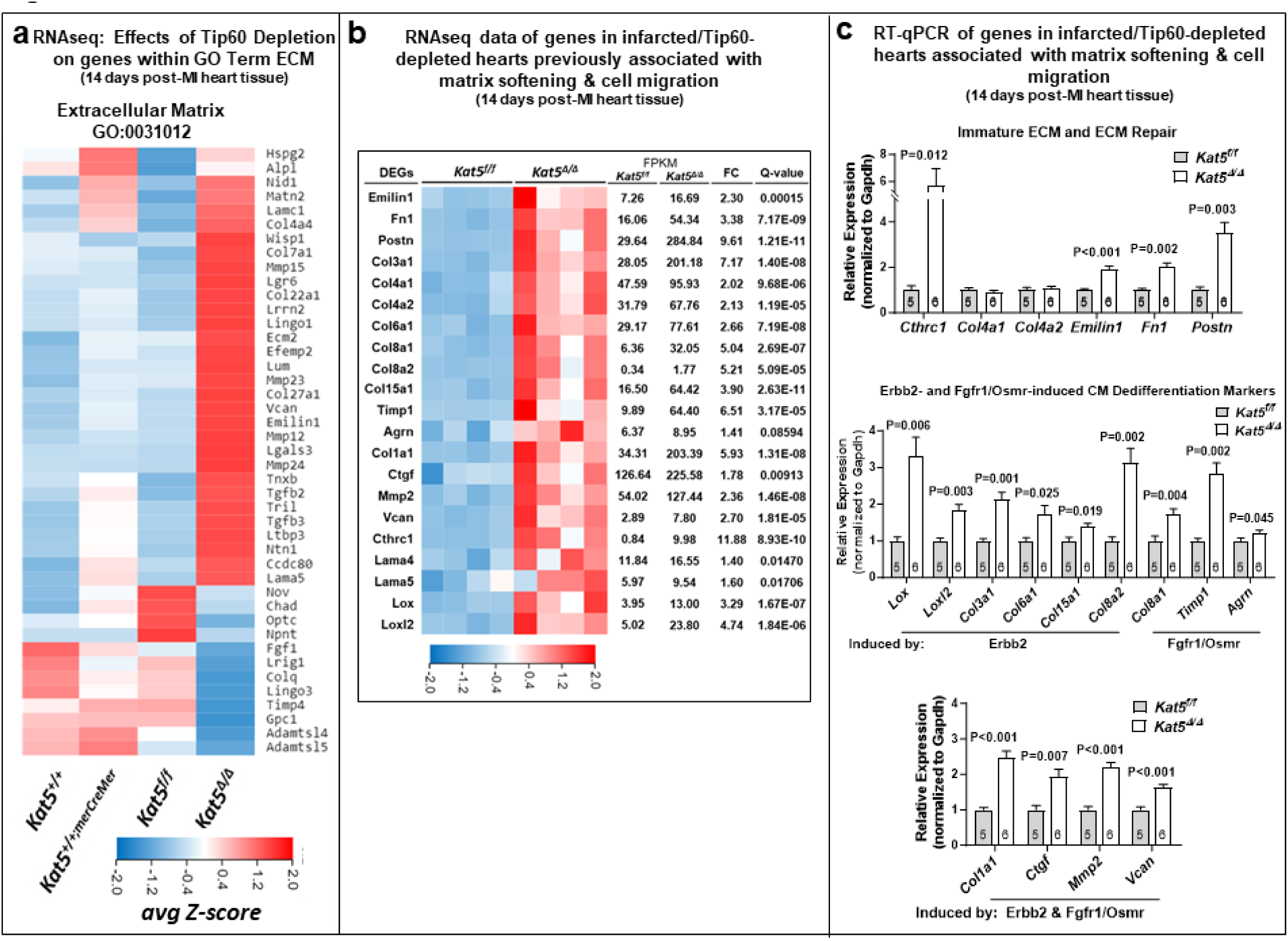
Gene expression indicating transition to soft connective tissue in infarcted/Tip60-depleted hearts. Panel. **a**, RNAseq data showing altered regulation of genes in the GO term ‘Extracellular Matrix’. **Panel b**, RNAseq data showing expression of ECM genes in infarcted/Tip60-depleted hearts. **Panel c**, RT-qPCR data showing expression ECM genes in Tip60-depleted hearts. Note that this includes genes previously shown by others to be increased by over-expressing Erbb2 or inducing Fgfr1/OsmR. *Kat5^f/f^* = control, *Kat5*^Δ^*^/^*^Δ^ = Tip60-depleted. In **a**, *Kat5^+/+^* and *Kat5^+/+;merCremer^* are controls to assess off-target effects of Cre. In **b**, each column represents a biological replicate. In **c**, each biological replicate (N) is a single heart each subjected to three technical replicates. Data are Mean ±SEM compared by unpaired, two-tailed Student’s t tests with Welch’s correction.

**Supplemental Figure 9.**
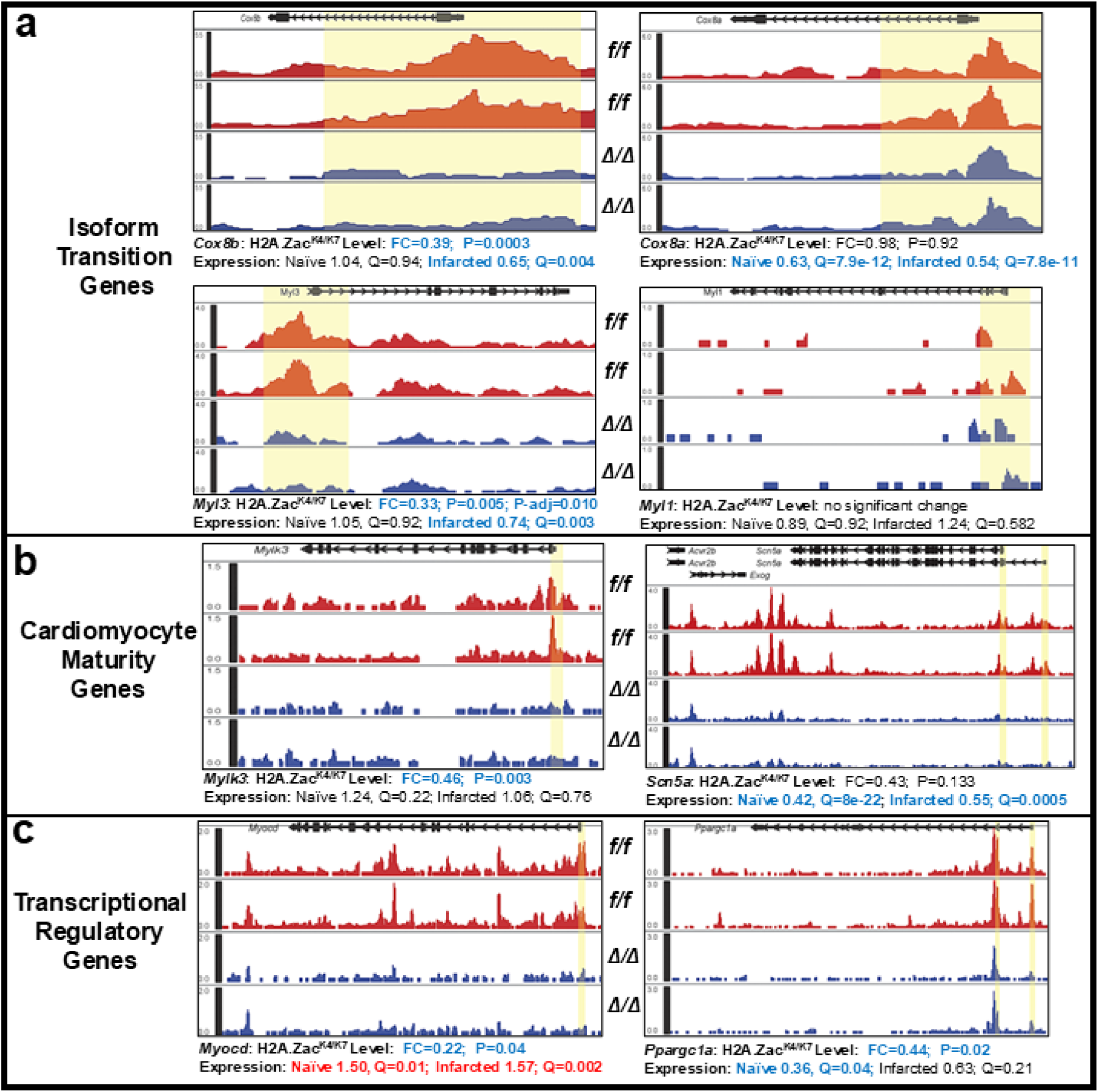
CUT&Tag localization of H2A.Zac^K4/K7^ in cardiac maturity genes (panels a-b) and in transcriptional regulatory genes (panel c). Unlike Figure 9 in which statistical significance of H2A.Zac^K4/K7^ depletion is based on p-adjust value, differences in this Figure are based on p-value. Yellow highlighting encloses ±1,000 bp around the TSS.

**Supplemental Figure 10.**
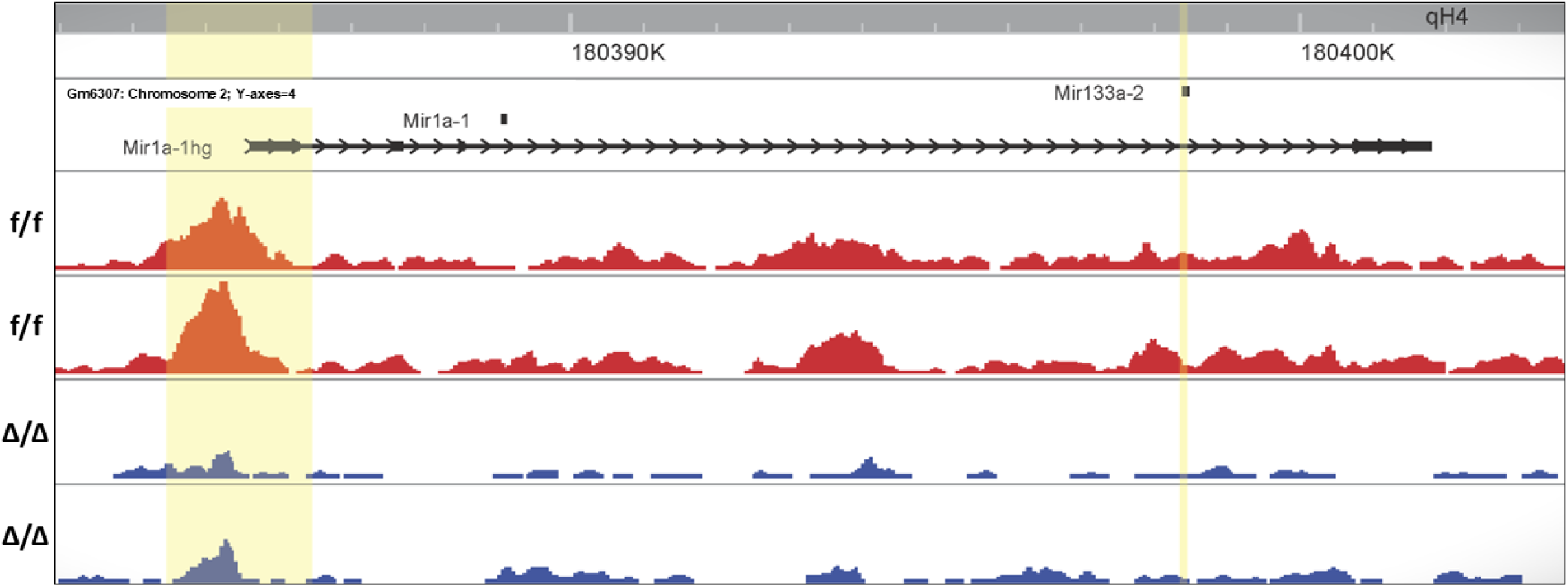
Tip60 KO-mediated reduction of H2A.Zac^K4/K7^ in the promoter/TSS locus of Mir1a-1hg (host gene; Gm6307), which encodes Mir1a-1 and Mir133a-2. These microRNAs are mapped as Gm6307, where they are separated by ∼9 kbp, transcribed as a bicistronic cluster. f/f = control; Δ/Δ = Tip60 KO (naïve); Fold-change = 0.341; P-value = 0.002; P-adjust value = 0.046.

**Supplemental Figure 11.**
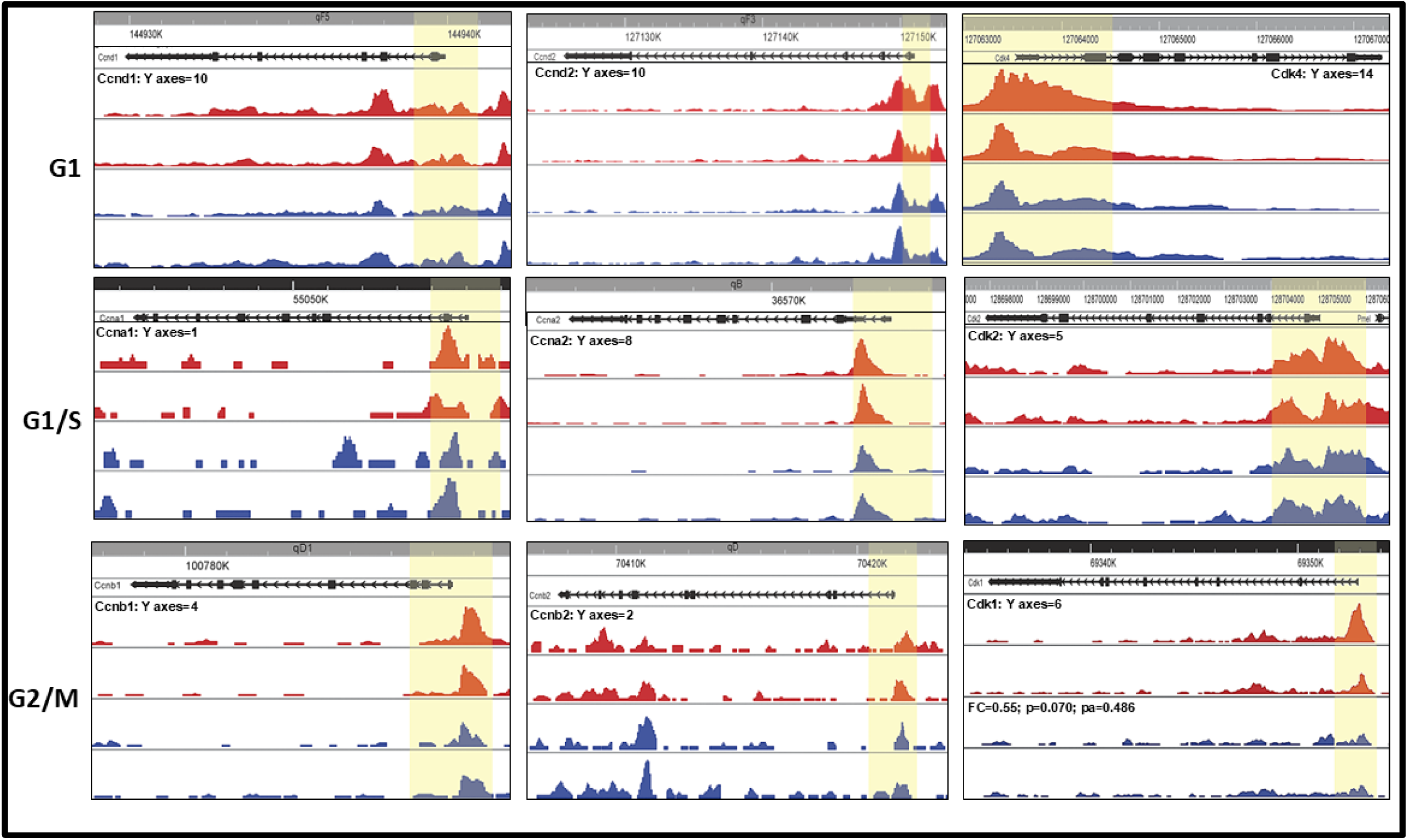
Tip60 KO does not affect H2A.Zac^K4/K7^ levels in cell-cycle genes. Among these and all other cell-cycle regulatory genes examined, the closest to exhibiting significant TSS/promoter locus depletion of H2A.Zac^K4/K7^ was *Cdk1* (lower right).

**Supplemental Figure 12.**
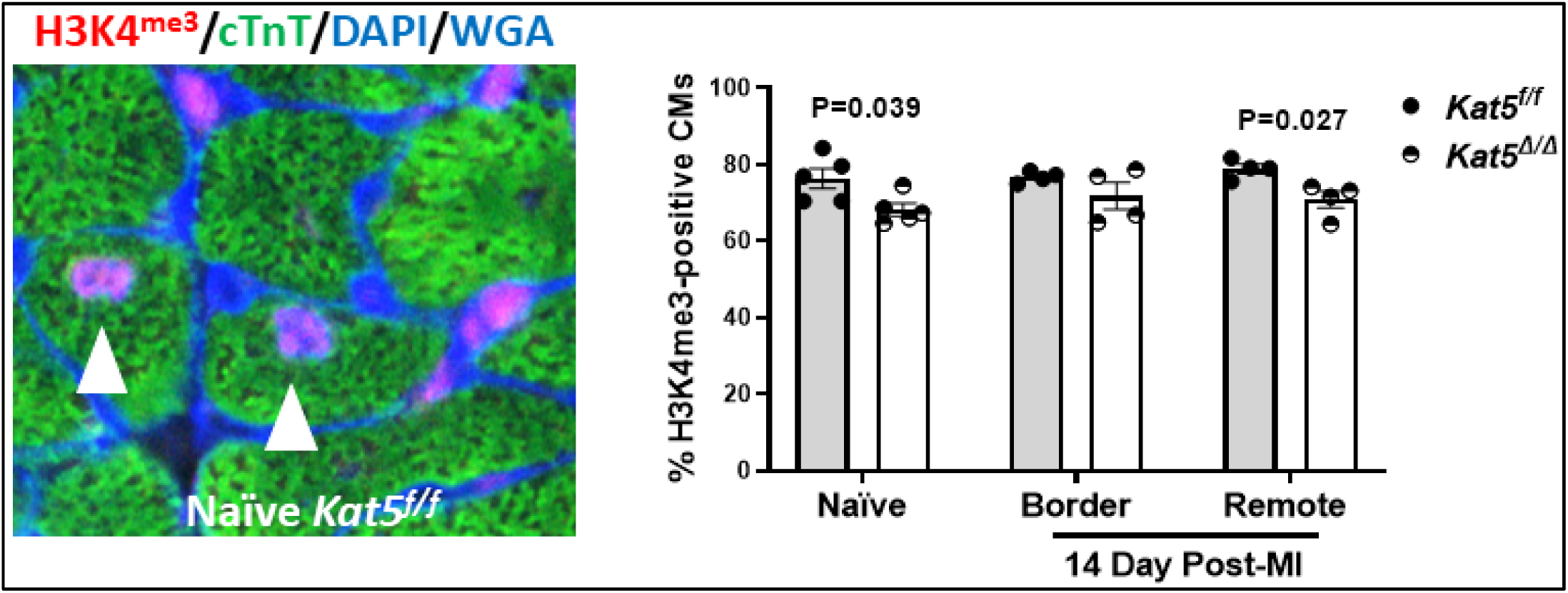
Subtly reduced levels of trimethylated histone H3 (H3K4^me3^) in Tip60-depleted Hearts. Panel. **a**, representative immunostained image showing H3K4^me3^-positive CMs in 13-week-old adult hearts from which Tip60 was depleted in CMs. Bars denote means ±SEM. *P<0.05 vs. *Kat5^f/f^* per unpaired Student’s t-test. Each point represents a biological replicate (i.e., individual mouse heart).

