## Supplemental Table 1 for "Loss of Tip60-dependent H2A.Z acetylation reprograms cardiomyocyte identity toward a regenerative state"

| Supplemental Table 1. Primers for PCR Genotyping | | | |
|---|---|---|---|
| Allele | Sequence (5'-3') and Working Conc. | Amplicon (bp) | Annealing ($^0$C) |
| *LoxP* in intron 2 | FWD GGAGGGAGTCAACGATCGCA 0.5 µM<br>REV AATGGGGGACCTACTCACCA 0.5 µM | 687 LoxP<br>586 WT | 61 |
| Cycling Details: 94 ºC 5 min, then 35 cycles of 94 ºC 30sec/61 ºC 45sec/72 ºC 45sec, then 72 ºC 10 mi | | | |
| *LoxP* in intron 11 | FWD GCACTCATCCAGGCTGTCC 0.5 µM<br>REV TCGGTTCTCAGAGACTAGC 0.5 µM | 655 LoxP<br>554 WT | 61 |
| Cycling Details: 94 ºC 5 min, then 35 cycles of 94 ºC 30sec/61 ºC 45sec/72 ºC 45sec, then 72 ºC 10 min | | | |
| *Myh6-Cre* transgene | FWD ATACCGGAGATCATGCAAGC 0.5 µM<br>REV AGGTGGACCTGATCATGGAG 0.5 µM | 440 | 54 |
| Cycling Details: 94 ºC 5 min, then 35 cycles of 94 ºC 30sec/54 ºC 45sec/72 ºC 45sec, then 72 ºC 10 min | | | |
| Internal control | FWD CTAGGCCACAGAATTGAAAGATCT 0.5 µM<br>REV GTAGGTGGAAATTCTAGCATCATCC 0.5 µM | 324 | 54 |
| Cycling Details: 94 ºC 5 min, then 35 cycles of 94 ºC 30sec/54 ºC 45sec/72 ºC 45sec, then 72 ºC 10 min | | | |

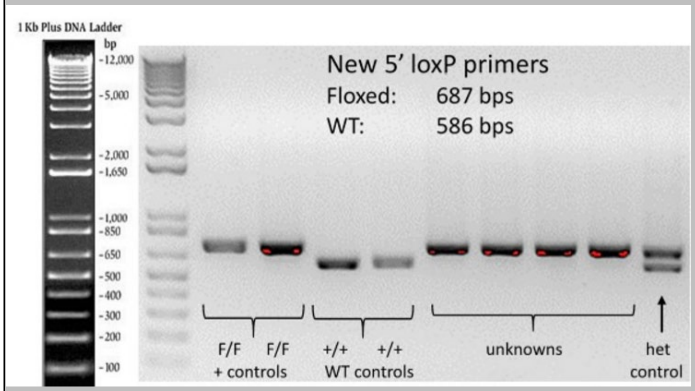

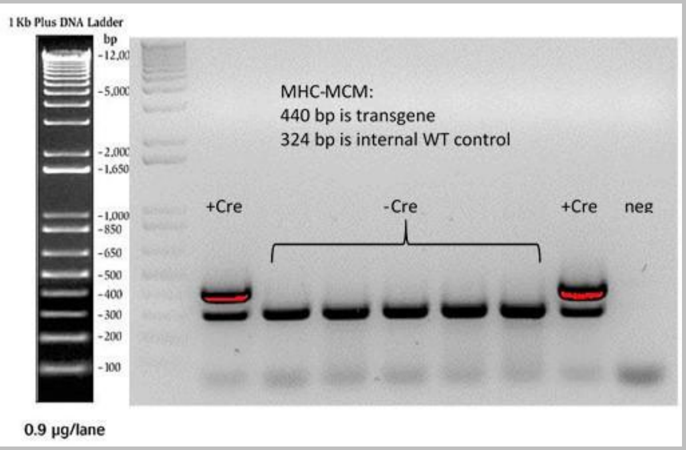
