## Supplemental Table 2 for "Loss of Tip60-dependent H2A.Z acetylation reprograms cardiomyocyte identity toward a regenerative state"

| Supplemental Table 2. Probes for Taqman qRT-PCR Gene Expression Analysis | |
|---|---|
| **Gene Target** | **Taqman Probe Kit** (Thermo-Fisher Catalog #) |
| Acadl | Mm01256456_m1 |
| Acadm | Mm01323360_g1 |
| Acadvl | Mm00444293_m1 |
| Acta2 | Mm00725412_s1 |
| Actn1 | Mm01304398_m1 |
| Agrn | Mm01264855-m1 |
| Col15a1 | Mm00456551_m1 |
| Col1a1 | Mm00801666_g1 |
| Col3a1 | Mm00802300_m1 |
| Col4a1 | Mm01210125_m1 |
| Col4a2 | Mm00802386_m1 |
| Col6a1 | Mm00487160_m1 |
| Col8a1 | Mm01344184_m1 |
| Col8a2 | Mm02344867_g1 |
| Cpt1b | Mm00487191_g1 |
| Cs | Mm00466043_m1 |
| Cthrc1 | Mm01163611_m1 |
| Cttn | Mm00514461_mH |
| Dab2 | Mm01307290_m1 |
| Ech1 | Mm00469322_m1 |
| Emilin | Mm0046244_m1 |
| Enol | Mm01619597_g1 |
| Fabp3 | Mm02342495_m1 |
| Fgf1 | Mm00438906_m1 |
| Fgfr1 | Mm00438930_m1 |
| Fgfr1 | Mm00438930_m1 |
| Fh1 | Mm01321349_m1 |
| Fn1 | Mm01256744_m1 |
| Gapdh | Mm99999915_g1 |
| Gja1 | Mm00439105_m1 |
| Hk1 | Mm00439344_m1 |
| Idh2 | Mm00612429_m1 |
| Kit | Mm0121916_m1 |
| Ldha | Mm01612132_g1 |
| Ldhb | Mm05874166_g1 |
| Lox | Mm00495386_m1 |
| Loxl2 | Mm00804740_m1 |
| Mapt | Mm00521988_m1 |
| Mdh2 | Mm00725890_s1 |
| Mef2c | Mm01340842_m1 |

| | |
|---|---|
| Mmp2 | Mm00439498_m1 |
| Myc | Mm00487804_m1 |
| Mylk4 | Mm01161239_m1 |
| Myocd | Mm00455051_m1 |
| Myom1 | Mm01237098_m1 |
| Nes | Mm00450205_m1 |
| Ogdh | Mm00803119_m1 |
| Osm | Mm01193966_m1 |
| Osmr | Mm01307326_m1 |
| Pdk3 | Mm00455220_m1 |
| Pfkl | Mm00435587_m1 |
| Pfkp | Mm00444792_m1 |
| Postn | Mm01284919_m1 |
| Runx1 | Mm01213404_m1 |
| Sdhb | Mm00458272_m1 |
| Shroom3 | Mm00497207_m1 |
| Slc16a3 | Mm00446102_m1 |
| Slc2a1 | Mm00441477_g1 |
| Slc2a4 | Mm00436615_m1 |
| Tbx20 | Mm00451515_m1 |
| Tgfb3 | Mm00436960_m1 |
| Tgfbr1 | Mm00436964_m1 |
| Tnni1 | Mm00502426_m1 |
| Tnni3 | Mm01330975_g1 |
| Vcan | Mm01283063_m1 |
