## Supplemental Table 3 for "Loss of Tip60-dependent H2A.Z acetylation reprograms cardiomyocyte identity toward a regenerative state"

### All Genes (56) Exhibiting Promoter/TSS Depletion of H2A.Zac$^{K4/K7}$ in Tip60 KO Hearts

**(details for genes shown in Fig. 10a heatmap)**

| **Note:** Nuclei for CUT&Tag were isolated from all cells contained in heart tissue samples, not from isolated CMs. Thus, only genes exhibiting >25% of total expression in CMs per Tabula Muris are considered CM genes. Genes shown in shaded rows are not considered CM genes. | | Effect of Tip60 KO on H2A.Zac$^{K4/K7}$ Level in Promoter/TSS Loci in Naïve Hearts CUT&Tag | | | Effect of Tip60 KO on Gene Expression in Infarcted Hearts RNAseq |
|---|---|---|---|---|---|
| **Gene** | **Protein** | **Fold-Change** | **P-value** | **P-adjust value** | **Fold-change/Q-value** |
| **Sh3kbp1**<br>% expression in CMs: 37<br>CM mean = 3.60 | Kinase-binding protein | **0.136** | 0.0002 | 0.012 | **1.24/0.030** |
| **Shroom3**<br>% expression in CMs: 74<br>CM mean = 1.66 | Shroom family member 3 | **0.164** | 9.1e-7 | 0.0003 | **1.7/0.000008** |
| **Irx4**<br>% expression in CMs: 100<br>CM mean = 0.41 | Iroquois homebox protein 4 | **0.165** | 0.0003 | 0.014 | 0.94/0.815 |
| **Asb14**<br>% expression in CMs: 99<br>CM mean = 2.62 | Ankyrin repeat/SOCS box-containing protein 14 | **0.169** | 9.11e-07 | 0.0003 | 0.98/0.941 |
| **Myh14**<br>% expression in CMs: 97<br>CM mean = 1.37 | Non-muscle myosin heavy chain II-C | **0.180** | 6.55e-06 | 0.001 | **0.62/0.002** |
| **Asb18**<br>% expression in CMs: 100<br>CM mean = 0.27 | Ankyrin repeat & SOCS box-containing protein 18 | **0.185** | 1.09e-5 | 0.002 | 0.85/0.559 |
| **Ces1d**<br>% expression in CMs: 68<br>CM mean = 3.85 | Carboxylesterase 1D | **0.189** | 0.0004 | 0.187 | **0.59/0.002** |
| **Abra**<br>% expression in CMs: 97<br>CM mean = 1.68 | Actin-binding rho- activating | **0.191** | 9.22e-06 | 0.0014 | 0.72/0.172 |
| **Tnni3**<br>% expression in CMs: 70<br>CM mean = 9.15 | Cardiac troponin I (cTnI) | **0.196** | 3.13e-06 | 0.0007 | **0.77/0.009** |
| **Dct**<br>% expression in CMs: 100<br>CM mean = 0.18 | Distal carboxyl-terminus (DCT) L-type Ca channel | **0.203** | 0.0016 | 0.045 | 0.95/0.95 |
| **Nkx2-5**<br>% expression in CMs: 96<br>CM mean = 1.81 | Homeobox protein Nkx2-5 | **0.221** | 7.61e-05 | 0.006 | 1.15/0.065 |
| **Gm3646**<br>% expression in CMs: 44<br>CM mean = 0.50 | lncRNA | **0.228** | 5.62e-07 | 0.00024 | 0.70/0.441 |
| **Rorc**<br>% expression in CMs: 87<br>CM mean = 0.16 | Retinoic acid receptor-related orphan receptor C | **0.228** | 5.85e-08 | 4.47e-05 | **0.42/0.00001** |
| **Trdn**<br>% expression in CMs: 95<br>CM mean = 5.02 | Triadin | **0.235** | 0.0003 | 0.015 | 0.87/0.271 |
| **Nlrp10**<br>% expression in CMs: 98<br>CM mean = 1.29 | Leucine-rich repeat containing 10 | **0.242** | 0.0012 | 0.037 | **0.67/0.017** |
| **Fsd2**<br>% expression in CMs: 94<br>CM mean = 4.02 | Fn type III & SPRY domain containing 2 | **0.243** | 6.80e-05 | 0.006 | 0.84/0.195 |
| **Gck**<br>% expression in CMs: 93<br>CM mean = 0.25 | Glucokinase | **0.246** | 3.07e-06 | 0.0007 | **0.30/0.0002** |
| **Acsm5**<br>% expression in CMs: 100<br>CM mean = 0.1 | Acyl-CoA synthetase medium-chain family member 5 | **0.250** | 0.0007 | 0.025 | **0.34/0.003** |
| **Lrrc52**<br>% expression in CMs: 100<br>CM mean = 0.1 | Leucine-rich repeat containing 52 | **0.253** | 1.25e-05 | 0.002 | 0.95/0.941 |

| | | | | | |
|---|---|---|---|---|---|
| **Unc45b**<br>% expression in CMs: 38<br>CM mean = 4.22 | UNC45B | **0.255** | 3.72e-08 | 3.46e-05 | **1.3/0.032** |
| **Gm20735**<br>(no expression data in Tabula Muris) | Semaphorin-4F | **0.259** | 0.0009 | 0.032 | no RNAseq expression data |
| **Ldb3**<br>% expression in CMs: 66<br>CM mean = 6.09 | Lim domain binding-3 | **0.260** | 3.11e-06 | 0.0007 | 1.2/0.079 |
| **Klhl40**<br>(no expression data in Tabula Muris) | Kelch-like protein 40 | **0.261** | 1.64e-08 | 1.98e-05 | 1.06/0.810 |
| **Ky**<br>% expression in CMs: 35<br>CM mean = 0.28 | Kyphoscoliosis Peptidase | **0.261** | 4.78e-05 | 0.004 | 0.69/0.067 |
| **Hhatl**<br>% expression in CMs: 99<br>CM mean = 3.85 | Hedgehog AT-like | **0.264** | 2.13e-05 | 0.002 | **0.32/3.15e-22** |
| **Klhdc7a**<br>% expression in CMs: 0<br>CM mean = 0.0 | Kelch domain-containing protein 7A | **0.267** | 9.40e-05 | 0.007 | 0.91/0.870 |
| **Mitf**<br>% expression in CMs: 38<br>CM mean = 1.82 | bHLH TF | **0.269** | 0.0001 | 0.007 | **0.56/0.0003** |
| **Homer2**<br>% expression in CMs: 89<br>CM mean = 1.81 | Homer scaffolding protein 2 | **0.271** | 0.0002 | **0.013** | **0.62/0.0002** |
| **Ckmt2**<br>% expression in CMs: 96<br>CM mean = 6.98 | Mitochondrial CK S-type | **0.273** | 3.79e-06 | 0.0008 | **0.41/9.55e-22** |
| **Fitm1**<br>% expression in CMs: 96<br>CM mean = 3.32 | Fat storage-inducing transmembrane protein 1 | **0.275** | 4.58e-08 | 3.76e-05 | **0.50/1.54e-9** |
| **Gm11213**<br>% expression in CMs: 0<br>CM mean = 0.0 | Calumenin | **0.279** | 0.0013 | 0.039 | no RNAseq expression data |
| **Gm5532**<br>(no expression data in Tabula Muris) | lncRNA | **0.285** | 8.87e-05 | 0.007 | no RNAseq expression data |
| **Tmem171**<br>% expression in CMs: 80<br>CM mean = 0.36 | Transmembrane protein 171 aka PRP2 | **0.299** | 0.0004 | 0.019 | 0.57/0.160 |
| **Hrc**<br>% expression in CMs: 65<br>CM mean = 4.64 | His-rich $Ca^{++}$-binding protein | **0.302** | 0.002 | 0.048 | **0.45/1.70e-8** |
| **Ckm**<br>% expression in CMs: 87<br>CM mean = 6.95 | Creatine kinase muscle Isoform | **0.306** | 9.8e-07 | 0.0003 | **0.59/6.08e-14** |
| **Ptgds**<br>% expression in CMs: 73<br>CM mean = 2.34 | Prostaglandin D2 synthase | **0.309** | 2.28e-05 | 0.003 | 0.67/0.402 |
| **Hrasls**<br>% expression in CMs: 99<br>CM mean = 1.51 | HRAS | **0.314** | 5.80e-07 | 0.0002 | **0.38/2.35e-15** |
| **Atp1b1**<br>% expression in CMs: 48<br>CM mean = 3.99 | Na+/K+ ATPase β-1 | **0.316** | 0.0003 | 0.017 | **0.74/0.0006** |
| **Ankrd23**<br>% expression in CMs: 84<br>CM mean = 2.20 | Ankyrin repeat: in I-band & ID | **0.317** | 6.58e-06 | 0.001 | **1.99/3.13e-7** |
| **Slc22a23**<br>% expression in CMs: 79<br>CM mean = 0.50 | Orphan transporter | **0.318** | 0.0001 | 0.009 | 0.84/0.400 |
| **Hfe2**<br>% expression in CMs: 95<br>CM mean = 3.86 | Hemojuvelin | **0.327** | 0.0002 | 0.011 | **0.34/8.12e-32** |
| **Rasgrp3**<br>% expression in CMs: 9<br>CM mean = 0.50 | Ras guanyl-releasing protein 3 | 0.329 | 0.0006 | 0.025 | 0.88/0.454 |

| | | | | | |
|---|---|---|---|---|---|
| **Gpt**<br>% expression in CMs: 69<br>CM mean = 0.93 | Glutamate-Pyruvate Transaminase | **0.331** | 4.93e-06 | 0.0009 | **0.49/0.00004** |
| **Tcap**<br>% expression in CMs: 83<br>CM mean = 6.90 | Telethonin | **0.331** | 0.002 | 0.044 | **1.49/0.022** |
| **Sema4b**<br>% expression in CMs: 16<br>CM mean = 0.22 | Semaphorin B | **0.337** | 2.59e-07 | 0.0001 | 1.00/0.985 |
| **Gm6307**<br>% expression in CMs: 100<br>CM mean = 0.19 | Genetic locus of microRNA miR-133a-2 | **0.341** | 0.002 | 0.046 | no RNAseq expression data |
| **Myo18b**<br>% expression in CMs: 89<br>CM mean = 2.97 | Myosin XVIIIb | **0.350** | 0.0001 | 0.009 | 0.91/0.616 |
| **Des**<br>% expression in CMs: 35<br>CM mean = 5.98 | Desmin | **0.356** | 0.0001 | 0.009 | **1.28/0.009** |
| **Rxrg**<br>% expression in CMs: 95<br>CM mean = 2.13 | Retinoid X receptor γ | **0.366** | 0.0008 | 0.028 | **0.59/5.15e-7** |
| **Cox8b**<br>% expression in CMs: 86<br>CM mean = 5.36 | Cytochrome c oxidase subunit 8b | **0.388** | 0.0003 | 0.015 | **0.65/0.004** |
| **Sox6**<br>% expression in CMs: 34<br>CM mean = 0.57 | Sox6 | **0.394** | 7.73e-06 | 0.001 | 0.76/0.229 |
| **Slc8a2**<br>% expression in CMs: 6<br>CM mean = 0.01 | NCX2 | **0.397** | 0.002 | 0.045 | 0.96/0.951 |
| **Prob1**<br>(no expression data in Tabula Muris) | Proline-rich basic protein 1 | **0.401** | 0.001 | 0.032 | 0.81/0.095 |
| **Speg**<br>% expression in CMs: 48<br>CM mean = 1.60 | Striated preferentially-expressed protein kinase | **0.456** | 3.35e-05 | 0.003 | **0.71/0.031** |
| **Car14**<br>% expression in CMs: 94<br>CM mean = 3.37 | Carbonic anhydrase XIV | **0.459** | 0.0007 | 0.028 | **0.28/2.82e-20** |
| **Crybb1**<br>% expression in CMs: 11<br>CM mean = 0.03 | Beta-crystallin B1 | 0.496 | 0.0005 | 0.020 | 0.41/0.030 |
| **Genes with increased H2A.Zac$^{K4/K7}$ in Promoter/TSS Locus** | | | | | |
| **Rab7b**<br>(no expression data in Tabula Muris) | Rab7b: role in mitophagy | **2.615** | 0.0016 | 0.046 | **1.85/0.023** |
| **Mhrt**<br>(no expression data in Tabula Muris) | Myheart | **3.410** | 1.59e-06 | 0.0004 | no RNAseq expression data |
