## Supplemental Table 4 for "Loss of Tip60-dependent H2A.Z acetylation reprograms cardiomyocyte identity toward a regenerative state"

### Effect of Tip60 Knockout on H2A.Zac$^{K4/K7}$ Levels & Gene Expression in CM Maturity Genes

| See Supplemental Table 3 for explanation of "% expression in CMs" value. | | Effect Tip60 KO on H2A.Zac$^{K4/K7}$ Level in Promoter/TSS of Naïve Hearts CUT&Tag | | Effect of Tip60 KO on Gene Expression in Infarcted Hearts RNAseq |
|---|---|---|---|---|
| **Gene** | **Protein** | **Fold-Change** | **P-value/P-adjust** | **Fold-change/Q-value** |
| **Cox8b**<br>% expression in CMs: 86<br>CM mean = 5.36 | Cytochrome C oxidase | **0.388** | **0.0003/0.015** | **0.65/0.004** |
| **Myl1**<br>% expression in CMs: 86<br>CM mean = 5.60 | Myosin light chain 1 (skeletal) | not affected | | **1.24/0.582** |
| **Myl3**<br>% expression in CMs: 79<br>CM mean = 4.34 | Myosin light chain 3 (cardiac) | **0.333** | **0.005/0.099**<br>**(P-adj=0.)** | **1.31/0.677** |
| **Mylk3**<br>% expression in CMs: 93<br>CM mean = 5.80 | Cardiac myosin light chain kinase | **0.255** | **0.023/0.257**<br>**0.003/0.065** | **1.06/0.755** |
| **Ppargc1α**<br>% expression in CMs: 90<br>CM mean = 2.86 | Peroxisome proliferator-activated receptor gamma coactivator 1-α | **0.440** | **0.020; 0.223** | **0.63/0.210** |
| **Ppargc1β**<br>% expression in CMs: 32<br>CM mean = 1.10 | Peroxisome proliferator-activated receptor gamma coactivator 1-β | not affected | | **1.42/0.060** |
| **Rxrα**<br>% expression in CMs: 22<br>CM mean = 0.83 | Retinoid X receptor α | not affected | | **0.81/0.067** |
| **Rxrβ**<br>% expression in CMs: 20<br>CM mean = 1.13 | Retinoid X receptor β | not affected | | **1.01/0.951** |
| **Rxrγ**<br>% expression in CMs: 95<br>CM mean = 2.13 | Retinoid X receptor γ | **0.367** | **0.0008/0.028** | **0.64/5.2e7** |
| **Rorc**<br>% expression in CMs: 87<br>CM mean = 1.53 | RAR-related orphan receptor C | **0.228** | **5.8e-8/4.5e-5** | **0.44/0.00001** |
| **Scn5a**<br>% expression in CMs: 97<br>CM mean = 2.25 | α-subunit (Nav1.5) of main cardiac voltage-gated sodium channel | **0.428** | **0.133/0.641** | **0.55/0.0005** |
| **Thra**<br>% expression in CMs: 20<br>CM mean = 1.55 | Thyroid hormone receptor α | not affected | | **0.59/9.26e-16** |
| **Tnni3k**<br>% expression in CMs: 99<br>CM mean = 2.85 | Troponin I-interacting kinase | **0.196** | **0.004/0.081** | **0.76/0.002** |
| **Cardiac Transcription Factor Genes** | | | | |
| **Gata4**<br>% expression in CMs: 22<br>CM mean = 2.31 | GATA binding protein 4 | not affected | | **1.25/0.036** |
| **Hand2**<br>% expression in CMs: 21<br>CM mean = 0.95 | Heart & Neural Crest 2 | not affected | | **1.16/0.042** |
| **Mef2a**<br>% expression in CMs: 15<br>CM mean = 2.42 | Myocyte enhancer factor 2A | not affected | | 0.95/0.748 |
| **Mef2c**<br>% expression in CMs: 11<br>CM mean = 2.07 | Myocyte enhancer factor 2C | not affected | | 1.04/0.889 |
| **Myocd**<br>% expression in CMs: 44<br>CM mean = 3.52 | Myocardin | **0.220** | **0.040/NA** | **1.57/0.002** |
| **Nkx2-5**<br>% expression in CMs: 96<br>CM mean = 1.81 | Nkx2-5<br>. | **0.221** | **7.6e-05/0.006** | 1.15/0.065 |
