## Supplemental Table 5 for "Loss of Tip60-dependent H2A.Z acetylation reprograms cardiomyocyte identity toward a regenerative state"

### Supplemental Table 5: Redifferentiation Markers

| Gene | Protein | Effect of Tip60 KO in Infarcted Hearts<br>RNAseq<br>Fold-change/Q-value |
|---|---|---|
| **Tumor Suppressors** | | |
| **Ptprv**<br>% expression in CMs: 0<br>CM mean = 0.00 | Protein Tyrosine Phosphatase Receptor Type V | 1.75/0.428 |
| **Ptprf**<br>% expression in CMs:7<br>CM mean = 0.27 | Protein Tyrosine Phosphatase, Receptor Type F | 1.25/0.288 |
| **Ptpn9**<br>% expression in CMs:5<br>CM mean = 0.30 | Protein Tyrosine Phosphatase Non-Receptor Type 9 | 1.13/0.424 |
| **Ptpn23**<br>% expression in CMs:17<br>CM mean = 0.43 | Protein Tyrosine Phosphatase Non-Receptor Type 23; maintains cardiac T-tubule. | 1.13/0.310 |
| **Phlda1**<br>% expression in CMs:17<br>CM mean = 0.74 | Pleckstrin homology-like domain family A member 1 | **1.52/0.004** |
| **Phlda3**<br>% expression in CMs:6<br>CM mean = 0.67 | Pleckstrin Homology-Like Domain Family A Member 3 | **1.83/0.014** |
| **Numbl**<br>% expression in CMs:3<br>CM mean = 0.11 | Numb-like protein | **1.77/0.0002** |
| **Dusp5**<br>% expression in CMs:28<br>CM mean = 0.38 | Dual Specificity Phosphatase 5; inhibits Erk1/2 | 1.37/0.301 |
| **Dusp6**<br>% expression in CMs:6<br>CM mean = 0.70 | Dual Specificity Phosphatase 6; inhibits Erk1/2 | 1.15/0.855 |
| **Spry1**<br>% expression in CMs:4<br>CM mean = 0.37 | Sprouty RTK Signaling Antagonist 1; inhibits receptor tyrosine kinase signaling; incl FGF | 1.24/0.212 |
| **Spry4**<br>% expression in CMs:5<br>CM mean = 0.38 | Sprouty RTK Signaling Antagonist 4; inhibits Mapk/Erk signaling; | 1.240.228 |
| **Nab2**<br>% expression in CMs:11<br>CM mean = 0.31 | Ngfi-a binding protein 2 | **1.63/0.004** |
| **Anti-Angiogenesis Factors** | | |
| **Col4a2**<br>% expression in CMs:10<br>CM mean = 1.93 | Type IV collagen α-2 chain; basement membrane component | **1.94/0.00001** |
| **Col4a3**<br>% expression in CMs:9<br>CM mean = 0.21 | Type IV collagen α-3 chain; basement membrane component; role kidney function; | 1.58/0.076 |
| **Col15a1**<br>% expression in CMs:5<br>CM mean = 0.28 | Type XV collagen α-chain; C-terminus inhibits angiogenesis; basement membrane component | **3.51/2.63e-11** |
| **Col18a1**<br>% expression in CMs:4<br>CM mean = 0.15 | Type XVIII collagen α-chain; C-terminus contains endostatin, which inhibits angiogenesis; basement membrane component | **1.51/0.022** |
| **Thbs1**<br>% expression in CMs:2<br>CM mean = 0.03 | Thrombospondin-1; ECM component; inhibits angiogenesis | **3.26/0.0001** |
| **Hippo Pathway Components** | | |
| **Mst1**<br>% expression in CMs: na<br>CM mean = 0.01 | Macrophage stimulating-1 aka HGF-like protein. tumor suppressor | 1.95/0.901 |
| **Lats1**<br>% expression in CMs:13<br>CM mean = 0.54 | Large Tumor Suppressor 1; phosphorylates & inhibits Yap/Taz; | 0.95/0.832 |
| **Lats2**<br>% expression in CMs:6<br>CM mean = 0.73 | Large Tumor Suppressor 2; phosphorylates & inhibits Yap/Taz; | **0.85/0.046** |
| **Mob1a**<br>% expression in CMs:6<br>CM mean = 0.53 | MOB kinase activator 1a; co-activates Lats1/2; tumor suppressor in context; | 1.13/0.471 |

| | | |
|---|---|---|
| **Mob1b**<br>% expression in CMs:21<br>CM mean = 0.45 | MOB kinase activator 1b; co-activates Lats1/2; tumor suppressor in context; | 1.21/0.514 |
| **Sav1**<br>% expression in CMs:10<br>CM mean = 0.24 | Salvador Family WW Domain-Containing Protein 1; acts as a scaffold in support of Mst1/2 kinases; tumor suppressor | 0.98/0.909 |
